# Bispecific Antibody Architecture and TNFRSF Target Selection Determine CD8^+^ T Cell Differentiation and Anti-tumour Immunity

**DOI:** 10.64898/2026.06.19.732642

**Authors:** Marcus A. Widdess, Lucy Wilkinson, Hannah J. Metcalfe, Anastasia Pakidi, H.T. Claude Chan, Jinny Kim, Tatyana Inzhelevskaya, Anna Turaj, Sean H Lim, Stephen Thirdborough, Stephen A Beers, Mark S Cragg, Aymen Al-Shamkhani

## Abstract

Tumour necrosis factor receptor superfamily (TNFRSF)-targeting bispecific antibodies (bsAb) enable tumour-localised T cell co-stimulation, yet how antibody architecture and receptor choice govern activity remains unclear. Here we define how bsAb format and TNFRSF target selection shape CD8^+^ T cell responses and anti-tumour immunity. Using B7-H3 as a tumour-associated antigen, we show that dual-bivalent 2×2 bsAb elicit maximal agonistic activity, whereas the corresponding monovalent 1×1 format is least active. A 2×1 format retaining TNFRSF bivalency but monovalent B7-H3 binding preserves substantial activity, identifying co-stimulatory receptor bivalency as a key determinant of efficacy. Across TNFRSF targets, 4-1BB drives the strongest cytotoxic CD8^+^ T cell differentiation and anti-tumour response. This superiority is conserved in human T cells and reproduced by cognate ligands, indicating that the observed functional hierarchy reflects receptor-intrinsic biology. Mechanistically, efficacy requires tumour-associated B7-H3 and T cell-dependent 4-1BB signalling. Together, these findings establish general principles linking antibody architecture and receptor biology to co-stimulatory bsAb efficacy and provide a framework for rational design leading to optimal therapeutics.

## Introduction

Cytotoxic CD8^+^ T cells are essential for tumour control, and insights into how cancers circumvent their surveillance have driven the development of therapies that mediate anti-tumour responses[1–3]. Consequently, immune checkpoint blockade has revolutionised cancer treatment, achieving durable responses and even cures in a subset of patients[4]. However, additional strategies are needed to broaden therapeutic options for patients who do not respond adequately to checkpoint blockade. Among the emerging approaches, targeting co-stimulatory receptors on T cells has shown considerable promise in pre-clinical studies[5–10]. Notably, co-stimulatory receptors within the tumour necrosis factor receptor superfamily (TNFRSF) constitute a major class of activating receptors that regulate T cell metabolism, proliferation, and survival, and support the differentiation of effector and memory T cells[11]. Importantly, engagement of these receptors has been shown to initiate transcriptional and metabolic programmes that synergise with checkpoint blockade by enhancing metabolic fitness and promoting robust CD8⁺ T cell expansion[12–16].

Despite the broad success of antibodies in biomedical applications, engineering agents to effectively agonise immune-stimulatory receptors has remained challenging[17]. First generation agonistic antibodies targeting TNFRSF members relied on co-engagement of Fcγ receptors (FcγR), particularly the inhibitory FcγRIIB, to induce sufficient antibody crosslinking, thereby facilitating efficient receptor clustering and activation of downstream TNFRSF signalling pathways[5, 10, 18]. However, clinical translation of these agonists has been largely hindered by limited efficacy, confounded by differences in the binding profiles of human and murine IgG isotypes to FcγR[17, 19]. To address these limitations, efforts have shifted toward developing bispecific antibodies (bsAb) that deliver agonistic signals independently of FcγR engagement. Thus, bsAb can deliver conditional agonism within the tumour microenvironment, provided that binding to the tumour-associated antigen allows efficient crosslinking of the TNFRSF target on T cells[20–30]. Although bsAb are now well established as a therapeutic modality, the relationship between molecular format and TNFRSF agonism remains unclear. As a result, design strategies are frequently determined by proprietary platform constraints rather than by systematic evaluation of alternative formats. Similarly, the impact of targeting different co-stimulatory receptors on the anti-tumour efficacy of bsAb is not well understood and rarely compared. Existing studies exhibit considerable variability in TNFRSF targets, antibody constructs, and experimental models, complicating the ability to reach a consensus on the most effective target and construct[20–30].

Here we conduct a systematic analysis of antibody format and immune receptor target selection to identify properties that determine bsAb functionality and anti-tumour efficacy. We find that both bsAb architecture and the choice of TNFRSF co-stimulatory target are critical factors influencing activity, underscoring the need for integrated optimisation of format and target in bsAb development.

## Results

### Valency and architecture modulate bsAb-mediated co-stimulation

We generated bsAb that activate the T cell co-stimulatory receptor 4-1BB (CD137) conditionally upon binding to tumour cells expressing the B7 homologue 3 (B7-H3; CD276). B7-H3 is an immunoglobulin superfamily protein overexpressed in multiple cancer types and has previously been targeted with antibodies, ADCs and chimeric antibody receptor T cells[31–33]. We evaluated different bsAb formats that varied in valency or spatial arrangement of their antigen-binding domains (Figure 1A). All bsAb were produced with a murine IgG1 (mIgG1) backbone incorporating the N297Q mutation, abrogating FcγR binding. We assessed the binding of the bsAb to their respective targets and found that bivalent constructs exhibited enhanced binding as determined by surface plasmon resonance (SPR) analysis (Supplementary Table 1), in line with their increased valency. Next, we evaluated their ability to co-stimulate OT-I T cell proliferation in the presence of either B7-H3-negative or B7-H3-positive tumour cells that were pulsed with ovalbumin 257-264 peptide (OVAp). To do this, we transduced MC38 tumour cells, which do not express endogenous B7-H3, with a construct encoding human B7-H3 (hB7-H3) or the equivalent empty vector to generate B7-H3-positive and control B7-H3-negative tumour cells, respectively (Supplementary Figure 1). Both the potency and maximal proliferative response, as assessed by CFSE dilution, improved with bsAb valency. The strongest effect was observed with the bsAb capable of bivalent binding to both targets (2×2), followed by the construct with bivalency for 4-1BB and monovalency for B7-H3 (2×1), while the bsAb with monovalent binding to both targets (1×1) was the least effective (Figure 1B). Interestingly, the 2×1 construct was significantly more active in the T cell co-stimulation assay compared to the 1×1 bsAb despite its weaker binding to B7-H3, suggesting that bivalency for 4-1BB is more critical than affinity for the tumour target (Figure 1B and Supplementary Table 1). Furthermore, none of the bsAb induced proliferation in the absence of B7-H3 and engagement of B7-H3 alone with a control 2×2 bsAb that only binds to B7-H3 failing to induce T cell proliferation (Figure 1B). Having established that dual bivalency per target enhances biological activity, we next explored an alternative 2×2 molecular format in which the anti-B7-H3 scFvs are appended to the C-terminus of the light chains (‘2×2 light’) instead of the heavy chains (Figure 1C). Two versions were generated: one with scFvs directly attached (as in the heavy chain 2×2 format), and another incorporating a flexible linker (G4S) between the C-terminus of the light chain and the scFv. Interestingly, although these ‘2×2 light’ bsAb bound 4-1BB comparably to the heavy-chain 2×2 bsAb, they both exhibited reduced activity in the T-cell co-stimulation assay (Figure 1C). While the construct lacking a linker showed negligible activity and weaker binding to B7-H3, incorporation of a flexible linker restored B7-H3 binding to levels comparable with the heavy-chain 2×2 format (Supplementary Table 1). These findings indicate that sub-optimal positioning of the anti-B7-H3 scFv can limit productive co-stimulatory signalling even in the presence of equivalent target binding. Based on these results, we selected the 2×2 heavy chain bsAb format as the optimal antibody architecture for subsequent investigation of different TNFRSF targets.

**Figure 1.**
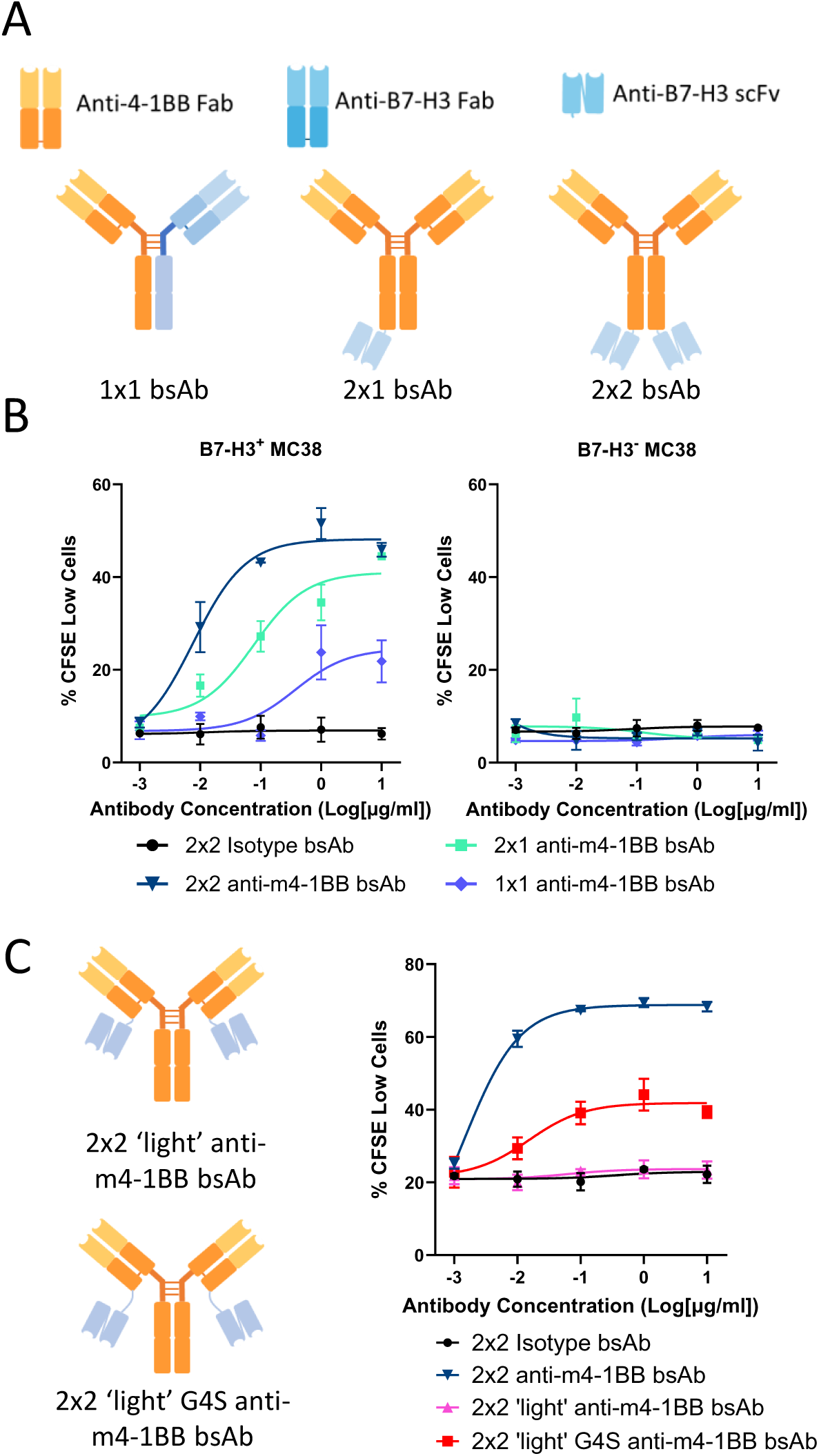
Anti-4-1BB bsAb format dictates CD8^+^ T cell co-stimulation potency. **a, c** Schematic representation of anti-m4-1BB x anti-B7-H3 bsAb formats used. **b** OT-I T cell proliferation determined by CFSE dilution 72 hours after stimulation with OVAp-pulsed MC38 B7-H3^+^ (left) or MC38 B7-H3-cells (right) and the specified bsAb. Data show mean ± SD and are representative of 3 independent experiments. **c** Comparison of the effects of different 2×2 anti-m4-1BB bsAb on OT-I T cell proliferation as determined by CFSE dilution 72 hours after stimulation with OVAp-pulsed MC38 B7-H3^+^ and the specified bsAb. Data show mean ± SD and are representative of 2 independent experiments.

### Distinct CD8^+^ T cell responses emerge from bsAb-targeting of different TNFRSF members

To evaluate other members of the TNFRSF, we produced three additional 2×2 bsAb (mIgG1; N297Q) targeting the co-stimulatory receptors CD27, GITR and OX40. The parental antibodies used to generate these bsAb were selected based on their previously reported agonistic activity[7, 9, 34]. Binding of these bsAb to B7-H3 and their respective TNFRSF targets was confirmed by SPR analysis (Supplementary Table 1). When comparing their effects on T cell proliferation, both the 4-1BB- and CD27 bsAb induced robust proliferative responses, with CFSE dilution observed in over 70% of T cells. The GITR-targeted bsAb showed a moderate effect, while the OX40 bsAb failed to elicit a proliferative response beyond the baseline level seen with the control bsAb (Figure 2A). The proliferative effects induced by the bsAb were diminished when MC38 cells lacked B7-H3, indicating that activity of the bsAb was conditional upon tethering by tumour cell expressed B7-H3 (Figure 2A). Examination of co-stimulatory receptor expression showed that whereas CD27, GITR and 4-1BB were expressed on OT-I T cells, OX40 was not upregulated under the conditions employed in this assay (Supplementary Figure 2). This finding is consistent with our previous observation that OX40 upregulation on CD8 T cells requires a higher antigen density compared to 4-1BB[9]. Examination of additional T cell functional traits revealed further differences in the response elicited by the various bsAb. Thus, secretion of IL-2 was demonstrable only with bsAb targeting 4-1BB and CD27, albeit more strongly with the anti-4-1BB bsAb (Figure 2B), while induction of CD25 and granzyme B were detected exclusively in the presence of the anti-4-1BB bsAb (Figure 2C and D).

**Figure 2:**
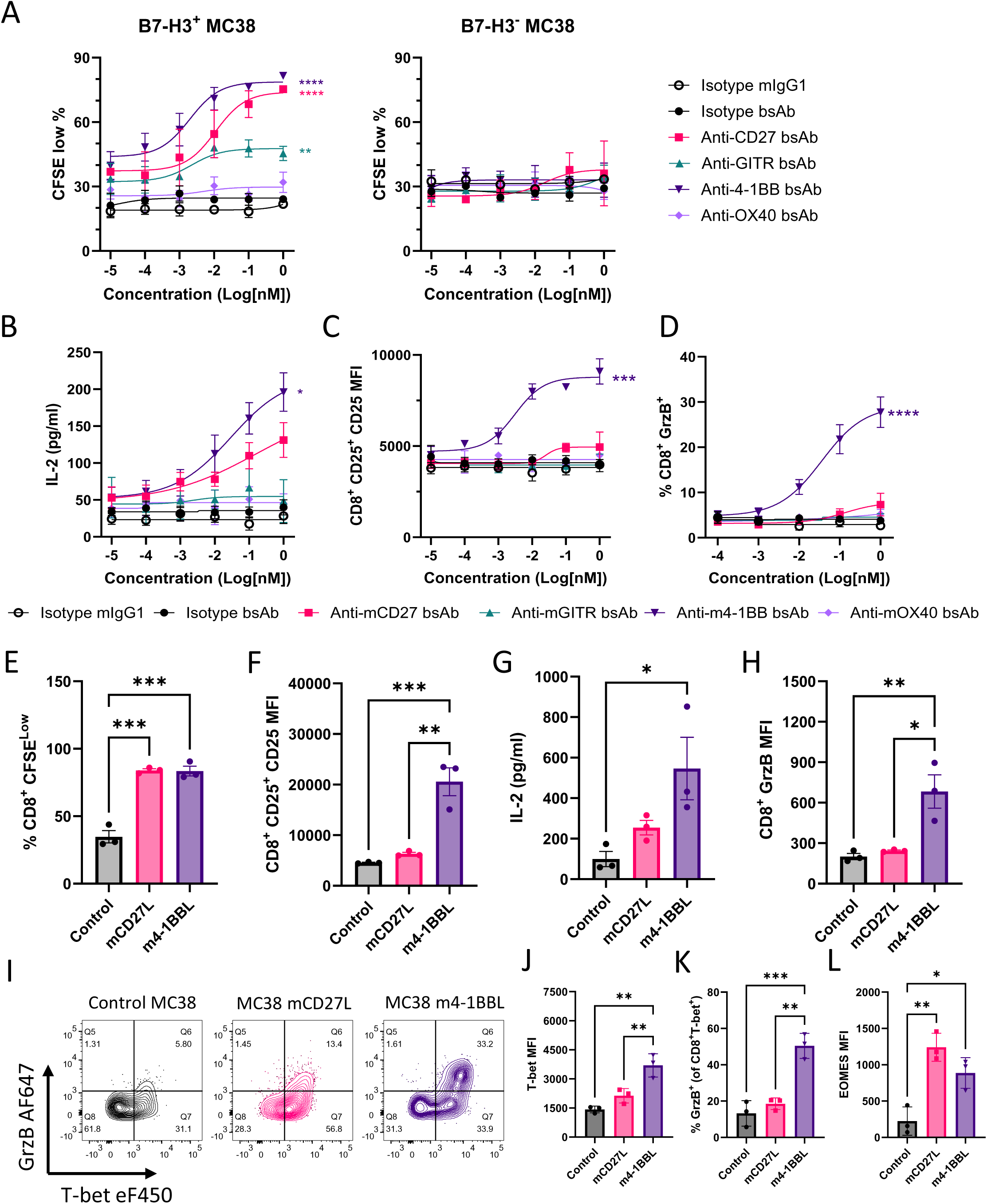
Distinct TNFRSF members drive divergent CD8⁺ T cell activation and differentiation programmes. **a** OT-I T cell proliferation determined by CFSE dilution 72 hours following stimulation with OVAp-pulsed hB7-H3^+^ MC38 (left) or hB7-H3^−^ MC38 (right) cells and the specified bsAb. **b-d**. **b** IL-2 concentration in supernatant, **c** CD25 expression and **d** granzyme B expression in co-cultures of OT-I T cells and OVAp-pulsed MC38 hB7-H3^+^ cells in the presence of the specified bsAb. Data (a-d) show mean ± SEM of 3 independent experiments with statistical significance determined using one-way ANOVA with multiple comparisons on AUC and significance shown between indicated antibody and isotype mIgG1. **e-h. e** CFSE dilution, **f** CD25 expression, **g** IL-2 concentration in supernatant and **h** granzyme B expression following OT-I splenocyte co-culture with OVAp-pulsed control MC38 cells or MC38 cells expressing CD27L or 4-1BBL. Data show mean ± SEM of 3 independent experiments, with statistical significance determined by one-way ANOVA. **i-l. i** Representative contour plots showing granzyme B and T-bet expression in OT-I T cells after stimulation with OVAp-pulsed MC38 cell lines. **j** T-bet MFI, **k** proportion of granzyme B^+^ cells in T-bet^+^ OT-I T cells after stimulation with OVAp-pulsed control MC38, MC38 mCD27L or MC38 m4-1BBL cells. Data show mean ± SEM of 3 independent experiments, with statistical significance determined by one-way ANOVA. *p<0.05, **p<0.01, ***p<0.001, ****p<0.0001.

To determine whether the distinct responses triggered by the anti-TNFRSF bsAb reflected the nature of the particular antibody components or inherent differences in receptor biology, we engineered MC38-hB7-H3 tumour cells to express the native ligands for CD27 or 4-1BB (Supplementary Figure 3). CD27L (CD70) and 4-1BBL co-stimulation induced a similar level of OT-I T cell proliferation, which was significantly greater than that seen with control MC38-B7-H3 cells (Figure 2E). In contrast, the production of IL-2, CD25 and granzyme B was primarily stimulated by 4-1BBL, mirroring the findings with the bsAb and suggesting that engagement of these two receptors triggers distinct effects on CD8^+^ T cell differentiation (Figure 2F-H). To further probe differences in CD27 and 4-1BB co-stimulation on CD8^+^ T cell differentiation, we examined the expression of the T-box transcription factors T-bet and eomesodermin (Eomes), which are known to play important roles in the formation and function of effector and memory CD8^+^ T cells[35]. 4-1BB co-stimulation was associated with increased T-bet expression and correspondingly higher granzyme B levels (Figure 2I–K). In contrast, although Eomes expression was upregulated following co-stimulation through both receptors, CD27 showed a trend toward higher expression compared with 4-1BB (Figure 2L). Collectively, these findings suggest that relative to CD27, 4-1BB co-stimulation preferentially drives differentiation toward an effector-like phenotype, highlighting distinct functional properties of these two co-stimulatory receptors.

Prompted by the contrasting effects of CD27 and 4-1BB co-stimulation on murine T cells, we next investigated whether human T cells responded similarly to bsAb targeting the corresponding human receptors. To minimise potential confounding effects arising from epitope specificity, we selected two antibody clones for each target that bind non-overlapping epitopes (CD27: Varlilumab and hCD27.15[36]); 4-1BB: Urelumab and Utomilumab[37]). All bsAbs were generated in a human IgG1 backbone containing PGLALA mutations to abrogate FcγR binding and Fc-mediated effector functions[38] and incorporated the same B7-H3-targeting scFv used in the murine bsAbs.

Human PBMCs were co-cultured with either control MC38-negative cells or hB7-H3^+^ MC38 cells together with a suboptimal concentration of anti-CD3. Co-cultures were treated with either isotype control B7-H3 only bsAb or co-stimulatory bsAbs, and CD8⁺ T cell proliferation, CD25 expression, and granzyme B expression assessed after 72 hours by flow cytometry. Minimal co-stimulatory activity was observed in the control MC38 co-cultures (Figure 3A-C upper panels). In contrast, in co-cultures containing hB7-H3^+^ MC38 cells, TNFRSF-targeting bsAbs significantly enhanced CD8⁺ T cell proliferation compared with control MC38 co-cultures (Figure 3A lower panel). Both anti-h4-1BB bsAb significantly increased CD25 expression relative to the isotype control (Figure 3B lower panel). Although the two anti-hCD27 bsAb promoted CD25 upregulation, the magnitude of this effect was substantially lower than that induced by 4-1BB co-stimulation (Figure 3B lower panel). We next assessed granzyme B expression specifically within divided CD8⁺ T cells, reasoning that these cells had received sufficient co-stimulatory signalling to undergo proliferation. Consistent with the CD25 data, both anti-h4-1BB bsAb markedly enhanced granzyme B expression, whereas anti-hCD27 bsAb had little effect relative to the isotype control (Figure 3C lower panel). Together, these findings mirrored the differential effects of CD27 and 4-1BB co-stimulation observed in murine CD8⁺ T cells, suggesting that these distinct co-stimulatory programs are conserved across species.

**Figure 3:**
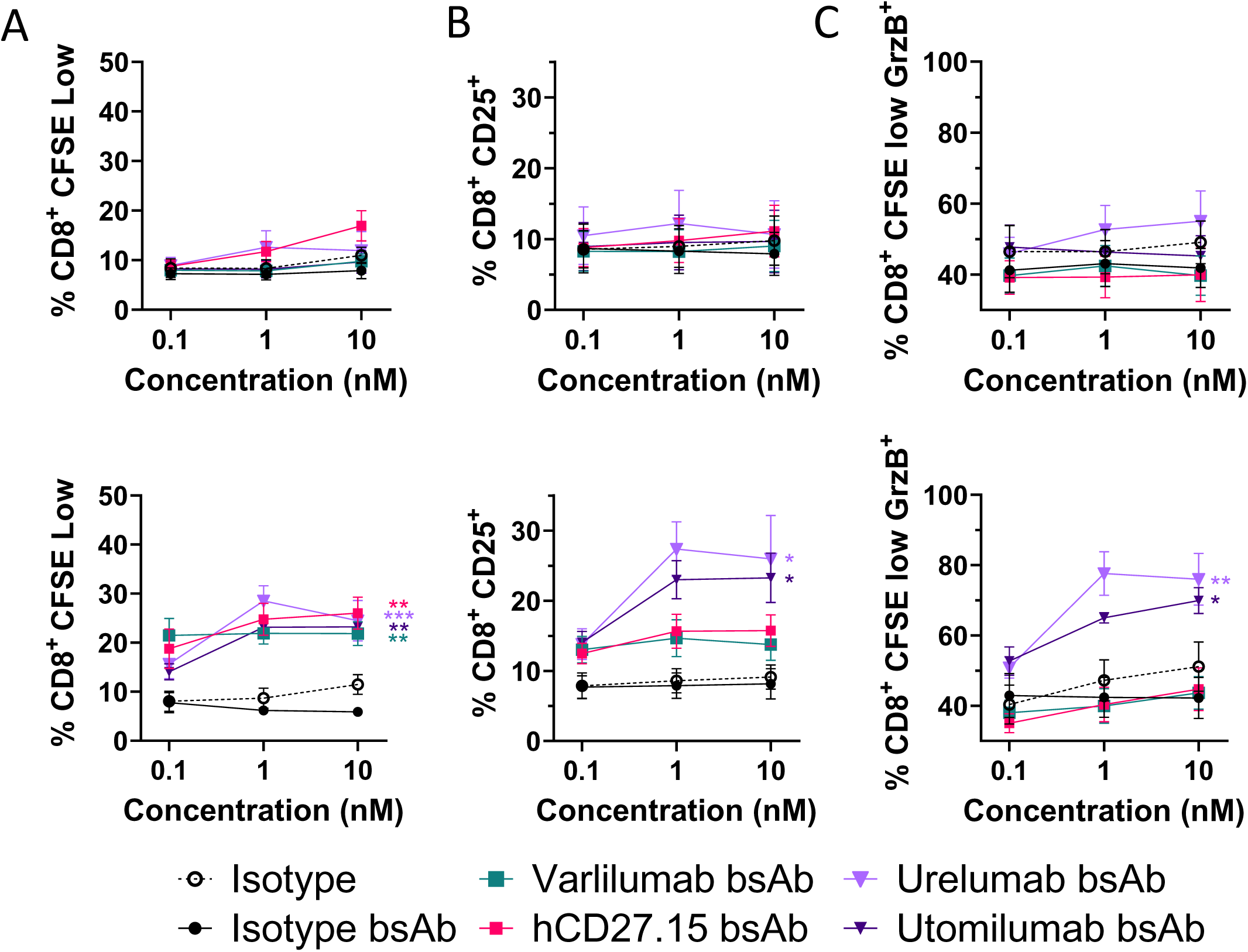
Contrasting effects of bsAb targeting CD27 or 4-1BB on human CD8^+^ T cell activation. **a-c** Human PBMCs co-cultured with control hB7-H3- MC38 (upper panels) or hB7-H3^+^ MC38 (lower panels) cells were stimulated with anti-CD3 and the specified antibodies for 72 hours. **a** CFSE dilution, **b** CD25 expression on CD8^+^ T cells, and **c** granzyme B expression on CD8^+^ CFSElow T cells. Data show mean ± SEM of 4 independent donors for CFSE dilution and mean ± SEM of 5 independent donors for CD25 and granzyme B expression. Statistical significance was determined using a one-way ANOVA on the 10 nM concentration with significance shown between the indicated antibody and isotype bsAb. *p<0.05, **p<0.01, ***p<0.001.

### Co-stimulatory target determines the anti-tumour efficacy of anti-TNFRSF bsAb

Given the distinct in vitro co-stimulatory effects of targeting different TNFRSF members, we proceeded to assess their potential to induce an anti-tumour immune response. Treatment of C57BL/6 mice bearing hB7-H3^+^ MC38 tumours with either anti-OX40 or anti-GITR bsAb was largely ineffective (Figure 4A and B). In contrast, anti-4-1BB bsAb treatment was highly protective in suppressing tumour growth with tumour regression observed in over 80% of the mice, while the anti-CD27 bsAb provided moderate protection with tumour regression seen in nearly half of the treated mice (Figure 4A and B). Moreover, the 4-1BB targeting bsAb also outperformed the CD27-targeting bsAb in the resistant 4T1 tumour model, providing a statistically significant delay in tumour growth whereas the anti-CD27 bsAb was ineffective (Supplementary Figure 4). To identify immune correlates of protective anti-tumour responses, we quantified tumour-infiltrating CD8^+^ T cells and assessed their differentiation into cytotoxic effector cells by measuring IFNγ, TNFα, and granzyme B expression. The 4-1BB- and CD27-targeting bsAb induced comparable increases in CD8^+^ T cell accumulation and the production of IFNγ and TNFα relative to controls, although only the effects of the 4-1BB-targeting bsAb reached statistical significance. In contrast, only 4-1BB targeting significantly increased the proportion of granzyme B-expressing CD8⁺ T cells (Figure 4D). This pattern mirrored our in vitro findings using either bsAb or the natural membrane-associated ligands (Figure 2). Thus, the ability of TNFRSF-targeting bsAb to promote effector-like CD8^+^T cell differentiation, as reflected by granzyme B expression, was more closely associated with anti-tumour efficacy than their capacity to drive T cell expansion.

**Figure 4:**
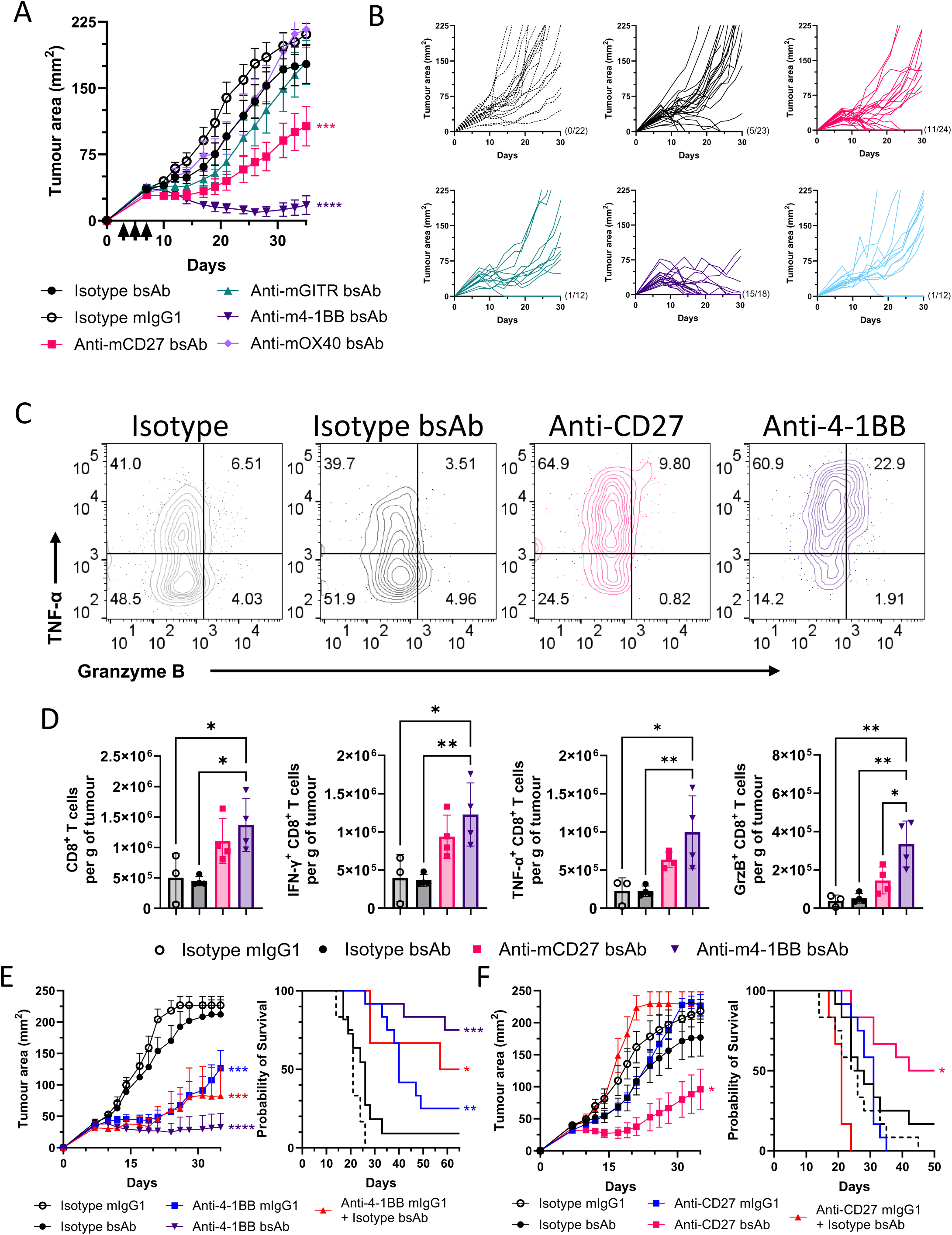
Co-stimulatory target determines the anti-tumour efficacy of anti-TNFRSF bsAbs. **a, b** Tumour area of mice inoculated with MC38 hB7-H3 tumour cells and treated with the specified antibodies on days 3, 5 and 7 (indicated by black arrows). **a** Data show mean tumour area ± SEM of 18-24 mice from 4 independent experiments and **b** individual tumour growth curves. Statistical significance was determined using one-way ANOVA with multiple comparisons on AUC. Significance shown between indicated antibody and isotype mIgG1. **c** Representative contour plots showing TNF-α and granzyme B expression in CD8^+^ T cells recovered from MC38 tumours treated with the indicated bsAb. **d** Number of CD8^+^ T cells (far left), IFN-γ^+^ CD8^+^ T cells (centre left), TNF-α^+^ CD8^+^ T cells (centre right) and granzyme B^+^ CD8^+^ T cells (far right) per gram of tumour following treatment with the indicated antibodies. Bar graphs indicate mean ± SEM of 4 mice with individual data points shown. Statistical significance established using one-way ANOVA with multiple comparisons. **e-f** MC38 hB7-H3 tumour growth curves (left) and survival plots of tumour-bearing mice (right) for the specified antibodies targeting either **e** 4-1BB or **f** CD27. Data show mean ± SEM of 11-12 individual mice from 2 independent experiments. Statistical significance for tumour area was determined using one-way ANOVA with multiple comparisons or by Log-rank (Mantel–Cox) test for survival. *p<0.05, **p<0.01, ***p<0.001, ****p<0.0001. Significance shown between indicated antibody and isotype mIgG1.

Next, we evaluated whether anti-CD27 and anti-4-1BB bsAb conferred superior anti-tumour immunity compared to their respective parental Fc competent mouse IgG1 antibodies. MC38-B7-H3 tumour-bearing mice were treated with the bsAb, the parental IgG antibodies alone, or the parental IgGs in combination with a control isotype x anti-B7-H3 bsAb. The anti-tumour activity of both the 4-1BB and CD27 bsAb surpassed that of their parental IgGs and was also greater than that achieved by combining the parental antibodies with the control anti-B7-H3 bsAb (Figure 4E and F). These findings demonstrate that the coordinated delivery of TNFRSF co-stimulation through a single tumour-targeted bsAb is more effective than providing the targeting and agonistic functions as separate molecules.

### Tumour cell B7-H3 and T cell-intrinsic 4-1BB signalling determine 4-1BB bsAb efficacy

Given that the anti-B7-H3 used in our bsAb recognises both human and murine B7-H3[33], we were able to investigate the relative contribution of tumour cell- and endothelial cell-associated B7-H3 to therapeutic activity. Since B7-H3 is expressed on the tumour cells and tumour endothelium[33], we examined whether expression on either compartment was sufficient for the activity of the anti-4-1BB x B7-H3 bsAb. First, we used immunofluorescence staining to validate that B7-H3 is expressed on the tumour endothelium in MC38 tumours. Expression of B7-H3 was co-localised with the endothelial marker CD31 in wild-type MC38 tumours, which contrasts with the widespread pattern of staining observed in the hB7-H3-transduced MC38 tumours (Figure 5A). Next, we investigated whether B7-H3 expression on the tumour cells was required for the anti-tumour response elicited by the 4-1BB bsAb. We treated control empty vector transduced MC38 or hB7-H3 transduced MC38 tumour bearing mice with either an isotype bsAb or the 4-1BB bsAb and observed tumour regression only in the 4-1BB bsAb treated mice bearing hB7-H3 transduced MC38 tumour cells (Figure 5B). These findings demonstrate that B7-H3 expression on endothelial cells alone is insufficient to mediate the anti-tumour activity of the 4-1BB bsAb.

**Figure 5:**
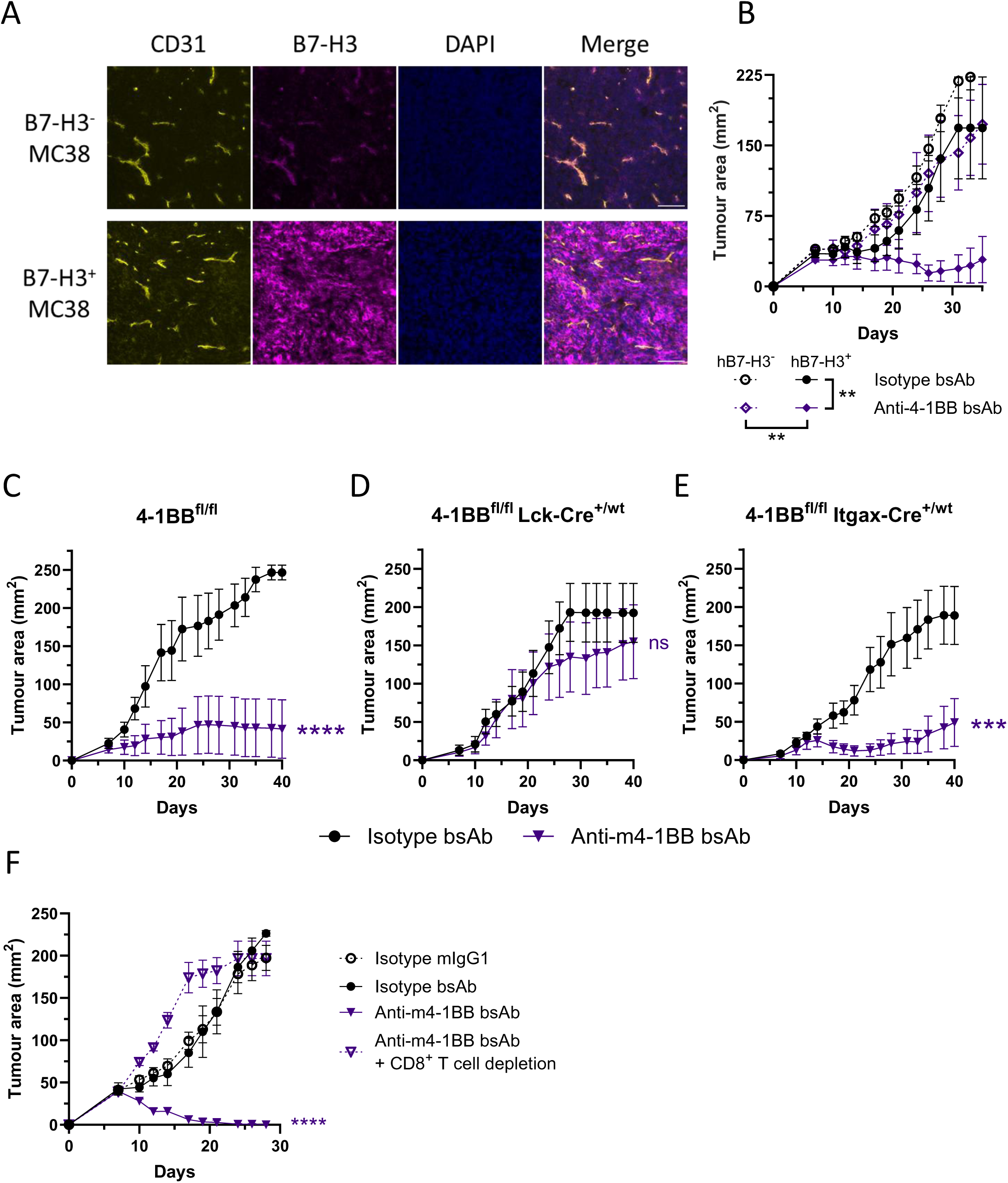
Anti-tumour activity of anti-4-1BB bsAb requires tumour cell B7-H3 and T cell 4-1BB expression. **a** Representative immunofluorescence images from B7-H3^−^ MC38 (upper) and hB7-H3^+^ MC38 (lower) showing CD31 (left, yellow), B7-H3 (centre left, magenta), DAPI (centre right, blue) and merged staining (right) showing CD31, B7-H3 and DAPI. Scale bar, 100 µm. **b** Tumour area of mice inoculated with MC38 B7-H3^−^ (hollow, dashed lines) or MC38 B7-H3^+^ (filled, solid lines) tumour cells and treated with the specified antibodies on days 3, 5 and 7. Data points show mean tumour area ± SEM of 6 mice per group. Statistical significance was determined by one-way ANOVA with multiple comparisons performed on AUC. **c-e** Tumour area in **c** 4-1BB^fl/fl^, **d** 4-1BB^fl/fl^ Lck-cre^+/wt^ and **e** 4-1BB^fl/fl^ Itgax-cre^+/wt^ C57BL/6 mice inoculated with MC38 hB7-H3 cells and treated with the specified antibodies on days 3, 5 and 7. Data points show mean tumour area ± SEM of 6-12 mice per group. **f** Anti-tumour activity of anti-4-1BB bsAb requires CD8^+^ T cells. Tumour area of mice inoculated with MC38 B7-H3 cells following treatment with the specified antibodies on days 3, 5 and 7. A group of anti-4-1BB bsAb treated mice were additionally injected with a depleting anti-CD8 antibody. Data points show mean ± SEM of 6 mice per group. Statistical significance by one-way ANOVA with multiple comparisons performed on AUC. *p<0.05, **p<0.01, ****p<0.0001. Significance shown between indicated treatment and isotype bsAb.

Next, we investigated the impact of B7-H3 density on the anti-tumour activity of the 4-1BB bsAb. We compared bsAb efficacy in mice bearing either control CT26 tumour cells, which endogenously express low levels of B7-H3, or CT26 tumour cells engineered to overexpress murine B7-H3 (Supplementary Figure 5A-B). The 4-1BB bsAb exhibited significant anti-tumour activity in both models, however, increased murine B7-H3 expression resulted in a higher proportion of mice undergoing complete tumour eradication (Supplementary Figure 5C-D), indicating that higher target antigen density enhances the therapeutic efficacy of 4-1BB-directed bsAb.

Although 4-1BB is classically described as a T cell co-stimulatory receptor, it is also expressed on dendritic cells (DC), where its engagement can induce pro-inflammatory cytokines including IL-12[39, 40], a key signal 3 cytokine that promotes effector CD8⁺ T cell differentiation and function[41, 42]. To determine whether the 4-1BB bsAb acted directly on CD8⁺ T cells or additionally through DC, we generated conditional 4-1BB knockout mice[5]. To generate mice with selective deletion of 4-1BB on T cells or DC, 4-1BB^fl/fl^ mice were crossed with either Lck-cre or Itgax-cre mice, respectively. Whereas the anti-4-1BB bsAb mediated robust anti-tumour activity in control 4-1BB^fl/fl^ mice (Figure 5C), this activity was abrogated in 4-1BB^fl/fl^ Lck-cre^+/wt^ mice (Figure 5D), demonstrating that intrinsic T cell co-stimulation by 4-1BB is essential for driving the therapeutic activity of the bsAb. In contrast, deletion of 4-1BB on DC had minimal impact on therapeutic efficacy, with the anti-4-1BB bsAb retaining robust anti-tumour activity in 4-1BB^fl/fl^ Itgax-cre^+/wt^ mice (Figure 5E). Consistent with these findings, depletion of CD8^+^ T cells in tumour bearing mice abrogated the ability of the 4-1BB bsAb to exert its anti-tumour activity (Figure 5F and Supplementary figure 6). Together, these findings indicate that the anti-4-1BB bsAb mediates its effects through CD8^+^ T cell co-stimulation rather than through activation of DC.

## Discussion

Despite compelling pre-clinical evidence supporting TNFRSF agonism as a strategy to enhance anti-tumour immunity, clinical translation has remained challenging[17]. Agonistic antibodies targeting receptors such as 4-1BB, CD27, OX40 and GITR have generally demonstrated limited efficacy as monotherapies, while dose-limiting toxicities, particularly hepatotoxicity with systemic 4-1BB agonism, have further constrained clinical development[17, 43]. A major obstacle has been the difficulty in achieving efficient receptor activation within tumours while avoiding widespread systemic activation. First-generation agonistic antibodies relied primarily on FcγR-mediated crosslinking to induce productive TNFRSF signalling[18], yet differences in FcγR expression and IgG biology between mice and humans likely contributed to the disconnect between pre-clinical efficacy and clinical activity[19]. Moreover, the relatively rare antibodies reported to exhibit FcγR-independent agonistic activity may promote widespread systemic receptor activation, thereby increasing the risk of dose-limiting toxicities[44–46]. Consequently, substantial effort has shifted toward the development of FcγR-independent bsAb designed to provide conditional agonism within the tumour microenvironment[47]. However, despite the rapid expansion of this field, there has been remarkably little systematic evaluation of how bsAb molecular architecture influences co-stimulatory function.

Here, we demonstrate that bsAb format is a critical determinant of agonistic activity. By directly comparing multiple architectures, we found that dual bivalency for both the tumour antigen and TNFRSF target produced the most potent co-stimulatory responses. Moreover, among the dual-bivalent constructs tested, appending the anti-B7-H3 scFv to the C-terminus of the heavy chain conferred superior activity compared with equivalent light-chain configurations, highlighting the importance of spatial organisation in bsAb function. Importantly, enhanced activity was not simply explained by improved binding to the tumour-associated antigen, as constructs with weaker B7-H3 binding but preserved 4-1BB bivalency (2×1) retained substantial activity. This observation also suggests that formats incorporating bivalency for the TNFRSF target but monovalent binding to the tumour antigen may provide an opportunity to increase tumour selectivity, as productive crosslinking would preferentially occur in tumours with high target antigen density while potentially limiting on-target off-tumour T cell activation in normal tissues expressing lower antigen levels. These findings indicate that bsAb architecture may need to be tailored according to the biological and expression characteristics of the tumour-associated target. Collectively, our findings support a model in which efficient TNFRSF receptor clustering is the dominant determinant of agonistic signalling. TNFRSF members are known to require receptor oligomerisation for productive recruitment of TRAF adaptor proteins and downstream NF-κB activation[10, 37, 48–50], and our data suggest that bsAb geometry critically influences the ability to nucleate these signalling assemblies. The superior activity of the 2×2 format therefore likely reflects a greater capacity to promote TNFRSF receptor clustering while simultaneously increasing the avidity of tumour-cell engagement. Bivalency for the TNFRSF target may enhance local receptor density and facilitate formation of signalling-competent receptor assemblies, whereas bivalent binding to the tumour-associated antigen may stabilise these interactions at the cell–cell interface by prolonging bsAb retention and reducing dissociation. These findings define key design principles that can guide the generation of candidates with optimal activity.

Beyond bsAb architecture, our data demonstrate that the choice of TNFRSF target fundamentally shapes both T cell differentiation and therapeutic efficacy. Although several TNFRSF members have been pursued clinically, comparative analyses between receptors have been limited by differences in experimental systems, antibody properties and tumour models. By systematically evaluating multiple receptors within a common bsAb format, we identify striking functional divergence between co-stimulatory pathways. Among the receptors tested, 4-1BB consistently elicited the strongest anti-tumour activity, whereas CD27 produced intermediate effects and GITR and OX40 exhibited minimal activity in our systems. Importantly, these differences were mirrored by distinct effects on CD8⁺ T cell differentiation. While both CD27 and 4-1BB promoted proliferation, only 4-1BB robustly induced CD25, granzyme B and T-bet expression, consistent with enhanced acquisition of an effector-like cytotoxic phenotype. Notably, engagement of CD27 and 4-1BB using their respective natural ligands recapitulated the differentiation programs induced by the corresponding bsAb, indicating that these effects reflect intrinsic differences in receptor biology rather than antibody-specific properties.

The lack of activity observed with OX40-targeting bsAb likely reflects the limited expression of OX40 on CD8⁺ T cells under our experimental conditions. Unlike CD27, GITR and 4-1BB, OX40 was not detectably upregulated on CD8⁺ T cells, consistent with previous reports that OX40 induction requires stronger or more sustained TCR signalling[9]. Consistent with this, whereas human OX40 is abundantly expressed on activated human CD4⁺ T cells, its expression on activated human CD8⁺ T cells is typically restricted to a relatively small subset of cells[51]. These findings highlight an important but often underappreciated consideration in co-stimulatory agonist development: receptor expression patterns, including both magnitude and temporal dynamics, may critically affect therapeutic responsiveness. In this context, CD27 is constitutively expressed on naïve T cells and available early during T cell priming and expansion, whereas 4-1BB is induced following activation and may preferentially reinforce highly activated effector T cell populations. Thus, differences in receptor expression dynamics, together with intrinsic differences in the magnitude or quality of downstream signalling[11, 52], likely contribute to the observed hierarchy of activity. Our findings therefore argue that TNFRSF receptors are not functionally interchangeable and that careful target selection is likely to be as important as bsAb format optimisation for achieving meaningful therapeutic activity.

Notably, the distinct functional programs induced by CD27 and 4-1BB were highly conserved between murine and human T cells. Across both species, 4-1BB co-stimulation preferentially promoted acquisition of cytotoxic effector traits, including granzyme B expression, whereas CD27 induced comparatively weaker effector differentiation despite robust proliferation. Importantly, these findings were reproduced using two independent bsAbs targeting non-overlapping epitopes for each receptor, supporting the generality of the observed effects. These observations strengthen the translational relevance of our findings and suggest that the superior anti-tumour efficacy of 4-1BB-targeting bsAb reflects conserved biology rather than antibody-specific differences.

Our study further demonstrates that the cellular context in which the tumour-associated antigen is expressed could be a critical determinant of effective conditional agonism. The finding that endothelial B7-H3 expression alone was insufficient to support therapeutic activity may reflect the inability of endothelial cells to provide tumour antigen-dependent signal 1 required for the initiation of CD8⁺ T cell activation. Consistent with an additional role for target antigen density in regulating bsAb activity, increasing B7-H3 expression on tumour cells further enhanced the efficacy of the 4-1BB bsAb, suggesting that antigen abundance directly influences the efficiency of receptor crosslinking and downstream signalling. Importantly, however, significant activity was still observed in tumours expressing relatively low endogenous levels of B7-H3, indicating that therapeutic activity can be achieved across a range of antigen densities. Together, these findings suggest that both the cellular context and abundance of tumour-associated antigen influence the activity of conditionally agonistic bsAb.

Finally, we demonstrate that the anti-tumour activity of the 4-1BB bsAb is primarily mediated through direct co-stimulation of CD8⁺ T cells rather than through activation of DC. In addition to its expression on activated T cells, 4-1BB is also expressed on DC, where its engagement can induce pro-inflammatory cytokines such as IL-12 that promote effector CD8⁺ T cell differentiation and function[39, 40]. However, our conditional deletion experiments demonstrated that therapeutic activity was critically dependent on T cell-intrinsic 4-1BB signalling, whereas deletion of 4-1BB from DC had minimal impact on anti-tumour activity. These findings indicate that the therapeutic effects of the bsAb are driven predominantly by direct co-stimulation of CD8⁺ T cells within the tumour microenvironment. Another notable finding was that both the CD27- and 4-1BB-targeting bsAb demonstrated greater anti-tumour efficacy than their respective parental Fc-competent antibodies, supporting the concept that tumour-targeted bsAb can deliver co-stimulatory agonism more effectively than conventional agonistic antibodies.

Collectively, our findings establish that both bsAb architecture and TNFRSF target selection are major determinants of therapeutic activity. More broadly, this work provides a mechanistic framework for the rational design of next-generation co-stimulatory bsAbs and suggests that optimal clinical activity will require integrated optimisation of receptor clustering, target antigen selection and co-stimulatory pathway biology.

## Methods

### Cell lines

CT26 colon carcinoma (ATCC, CRL-2638) and 4T1 (ATCC, CRL-2539) cells were grown in Roswell Park Memorial Institute (RPMI) medium 1640 (Gibco, 11875093), supplemented with 10% v/v foetal bovine serum (FBS, Sigma, F9665), 2 mM L-Glutamine (Gibco, 25030081), 1 mM Sodium Pyruvate (Gibco, 11360070), 100 U/ml Penicillin (Sigma, P4333-100ML), 100 μg/ml Streptomycin (Sigma, P4333) (termed complete RPMI; cRPMI). MC38 cells (kind gift from Dr Rienk Offringa, German Cancer Research Center, Heidelberg), Phoenix Ecotropic (Phoenix Eco, ATCC, CRL-3214) and A549 (ATCC, CCL-185) were grown in complete Dulbecco’s Modified Eagle Medium (cDMEM, Gibco), DMEM supplemented with 10% v/v FBS, 2 mM L-Glutamine, 1 mM Sodium Pyruvate, 100 U/ml Penicillin, 100 μg/ml Streptomycin.

### Retroviral transduction

MC38 and CT26 cells were stably transduced using the phoenix ecotropic retrovirus system[53]. Phoenix Eco cells were plated at 0.15 × 10^6^ cells per well in a 6 well plate on day -3. On day 0, Phoenix Eco cells were transfected with 4 µg of pCLEco (Addgene, 12371[53]) and 4 µg of the retroviral vector (pMP71 for CD27L and 4-1BBL and pMSCV-IRES-Thy1.1 for B7-H3; backbone plasmids from Professor Hans Strauss and Dr Thomas Mitchell), using the FuGENE HD transfection system. In brief, DNA was mixed in 135 µL antibiotic-free DMEM prior to adding 15 µL FuGENE HD transfection reagent (Promega, E2311). The transfection mixture was incubated at RT for 15 mins before adding to the Phoenix Eco cells in a dropwise manner. After 2 days, supernatant was collected and contaminating cells removed by centrifugation and 0.45 µm filtering. MC38 or CT26 cells were plated at 1×10^5^ cells per well of a 6 well plate on day 0. 2 ml of supernatant containing viral particles was added to the target cells on day 2 with 4 µg mL^−1^ of polybrene (Sigma, H9268). The plate was then centrifuged at 650 g for 90 mins at 32 °C. After 48 hours; cells were sorted via FACS, for GFP, or magnetic separation, for Thy1.1 expression.

### Antibodies

The sequences of the following antibodies were derived in-house from hybridomas: AT124-1 (anti-mouse CD27), LOB12.0 (anti-mouse 4-1BB), DTA-1 (anti-mouse GITR), MRC OX86 (anti-mouse OX40), AT171-2 (isotype control). Antibody sequences to human targets were derived from patents: anti-B7-H3 (m8524; US10604582), Utomilumab (WO2015/119923), Urelumab (WO2010/042433A1) Varlilumab (US9169325), hCD27.15 (US Patent 9527916). Recombinant antibodies were produced as previously described[10]. To produce bsAb with a 2×2 format, the anti-B7-H3 (m8524) scFv was genetically linked to the C-terminus of either the heavy or light chain sequence of anti-TNFRSF antibodies. To generate the 2×1 bsAb, complementary heterodimerisation mutations[54] were introduced into the CH3 domains of the anti-mouse 4-1BB heavy chains, and an anti-B7-H3 scFv was genetically fused to the C-terminus of one heavy chain. The 1×1 anti-mouse 4-1BB x B7-H3 bsAb was produced using the Fab arm exchange method as previously described[55] and heterodimerisation confirmed by hydrophobicity interaction chromatography (TSK Butyl-NPR, Tosoh Bioscience). BsAb were produced with either mouse IgG1 or human IgG1 backbones comprising the Fc disabling mutations NQ or PGLALA, respectively. Routinely antibody preparations contained < 1% aggregates as assessed by size exclusion chromatography (Zorbax GF-250, Agilent). All antibody preparations were endotoxin low (<5 EU endotoxin/mg) as determined using the Endosafe-PTS portable test system (Charles River Laboratories).

### Mice

C57BL/6J (Charles River UK, Strain code: 027), Balb/c (Strain code: 028, Charles River UK), and OT-I transgenic (Tg) mice (Charles River France, strain code: 642) were purchased from Charles River and stock colonies maintained by the University of Southampton Biomedical Research Facility. 4-1BB^fl/fl^ C57BL/6 mice were purchased as Tnfrsf9^tm1a(EUCOMM)Wtsi^ C57BL/6NTac mice from the International Mouse Phenotyping Consortium (IMPC, EPD0177_2_B12) and produced as previously described[5] and stock colonies maintained by the University of Southampton Biomedical Research Facility. Itgax^WT/Cre^ C57BL/6 mice (The Jackson Laboratories, #008068) and Lck^WT/Cre^ C57BL/6 mice (The Jackson Laboratories, #012837) were purchased from Jackson immunoresearch and stock colonies maintained by the University of Southampton Biomedical Research Facility. All mice were randomly assigned into experimental groups, with experimental and control animals co-housed. Mice were maintained on a 12-h light and dark cycle, an ambient temperature of 20–24 °C, 55% humidity ± 15%, with food and water ad libitum. Mice were kept under specific pathogen-free (SPF) conditions. Daily checks were performed to ensure mice remained healthy, and environmental enrichment was provided. Mice were euthanised by CO_2_ inhalation or cervical dislocation. All experiments were conducted following University of Southampton ethical approval and in accordance with the Animals (Scientific Procedures) Act 1986 as set out in PPL: P4D9C89EA and PIL: I66C5D543.

### Tumour models

Control MC38 cells transduced with empty vector or MC38 hB7-H3 cells (5×10^5^ cells) were subcutaneously injected into either the hind leg of C57BL/6 mice, 4-1BB^fl/fl^ Itgax^WT/Cre^ mice or 4-1BB^fl/fl^ Lck^WT/Cre^ mice on day 0. For immunotherapy experiments, mice were intraperitoneally injected with 500 pmole (75 µg for mIgG1, 100 µg for bsAb) of either isotype mIgG1, isotype bsAb, anti-GITR bsAb, anti-CD27 bsAb, anti-4-1BB bsAb or anti-OX40 bsAb on days 3, 5 and 7 (as specified in figure). Tumour growth was monitored every 2–3 days by calliper, and mice euthanised when cross-sectional area reached 225 mm^2^. Control CT26 cells transduced with empty vector or CT26 mB7-H3 cells (5×10^5^ cells) were subcutaneously injected into the hind leg of Balb/c mice on day 0. For immunotherapy experiments, mice were intraperitoneally injected with 500 pmole (75 µg for mIgG1, 100 µg for bsAb) of either isotype mIgG1, isotype bsAb, or anti-4-1BB bsAb on days 5, 7 and 9 (as specified in figure). Tumour growth was monitored every 2–3 days by calliper, and mice euthanised when cross-sectional area reached 225 mm^2^. 4T1 cells (2.5×10^4^ cells) were subcutaneously injected into the mammary fat pad of Balb/c mice on day 0. For immunotherapy experiments, mice were intraperitoneally injected with 500 pmole (75 µg for mIgG1, 100 µg for bsAb) of either isotype mIgG1, isotype bsAb, or anti-4-1BB bsAb on days 3, 5 and 7 (as specified in figure). Tumour growth was monitored every 2-3 days by calliper, and mice euthanised when cross-sectional area reached 225 mm^2^. For all tumour experiments, early termination was carried out if mouse welfare was deemed compromised, in accordance with the guidelines proposed by Foltz and Cullere[56].

### CD8^+^ T cell depletion in MC38 hB7-H3 tumour model

On day -2 and day 1, mice were intraperitoneally injected with 100 µg of anti-CD8 (clone: YTS169) mIgG2a (produced in house), to deplete CD8^+^ T cells. Depletion was confirmed by flow cytometric analysis of blood samples taken on days -1, 2, 8 and 14. MC38 hB7-H3 cells (5×10^5^) were subcutaneously injected into the hind leg of C57BL/6 mice on day 0. Mice were then intraperitoneally injected with 500 pmole (75 µg for mIgG1, 100 µg for bsAb) of either isotype mIgG1, isotype bsAb, or anti-4-1BB bsAb on days 3, 5 and 7 (as specified in figure). Tumour growth was monitored every 2–3 days by calliper, and mice euthanised when cross-sectional area reached 225 mm^2^.

### Ex Vivo restimulation of tumour infiltrating lymphocytes (TILs)

MC38 hB7-H3 cells (5×10^5^) were subcutaneously injected into the hind leg of C57BL/6 mice before they were treated with 500 pmole (75 µg for mIgG1, 100 µg for bsAb) of either isotype mIgG1, isotype bsAb, or anti-4-1BB bsAb on days 3, 5 and 7. Mice were euthanized on day 10 and the tumours were harvested and then injected with RPMI media containing 0.5 Wunsch Unit/ml Liberase DL (Roche) and 50 μg/ml DNase I (Roche) before incubating at RT for 5 minutes. Tumours were then dissected into 2 mm x 2 mm pieces and incubated for 20 mins at 37 °C under agitation. The digestion was quenched with RPMI containing 10% FBS, 10 mM EDTA and 20mM HEPES at room temperature (RT) for 10 mins. Tumours were then passed through a 100 μm cell strainer and the strainer rinsed with PBS to ensure the remaining cells were collected. The cell suspension was washed twice in PBS. TILs were resuspended at 1×10^6^ cells/ml in RPMI and restimulated with 20 ng/ml phorbol 12-myristate 13-acetate (PMA) and 1µg/ml ionomycin for 4 hours in the presence of Brefeldin A (BD Biosciences, 555029) and Monensin (BD Biosciences, 554724). The cells were then collected, washed twice with PBS and stained for flow cytometry. Gating strategy shown in Supplementary Figure 7.

### Immunofluorescence

C57BL/6 mice were subcutaneously injected with control MC38 or MC38 hB7-H3 cells (5×10^5^) on day 0. Tumours were harvested on day 12 and then frozen in optimal cutting temperature compound (OCT, CellPath), with the moulds incubated in a petri dish of isopentane (VWR chemicals) cooled by dry ice. Once frozen, tumour blocks were then removed from the moulds and sectioned using a Leica CM1860 UV cryostat (Leica). Sections were then placed onto microscope slides and incubated at RT overnight to ensure attachment. For immunofluorescence staining, sections were fixed in 100 % acetone (VWR Chemicals) for 10 mins at RT and then air dried for 5 mins to remove excess acetone. Sections were marked using a hydrophobic barrier pen before washing three times with PBS-0.05 % v/v Tween 20 (PBS-T). Sections were then blocked with PBS-T + 2.5 % v/v normal goat serum (VectorLabs, S-1000) for 30 mins at RT. Next, sections were stained using anti-CD31 (1:200, Bio-Rad, MCA2388) and anti-B7-H3 (1.4 μg/ml, Abcam, ab134161) in 100 μl PBS-T for 1.5 hours at RT in the dark. Antibody was removed and the sections washed 3 times with PBS-T. The sections were then stained with goat anti-rat IgG – AF568 (1:500, ThermoFisher, A-11077) and goat anti-rabbit IgG – AF488 (1:500, ThermoFisher, A-11006) in 100 μl PBS-T for 1.5 hours at RT in the dark. The secondary antibody was then removed, and the cells washed twice in PBS-T and once in PBS. VECTASHIELD HardSet antifade mounting medium (Vector, H-1400-10) was then added to the sections before coverslips placed on top and left to set overnight. The slides were then imaged using an Olympus CKX41 microscope and cellSens Standard (Olympus, version 2.3) software.

### OT-I Tg CD8^+^ T cell proliferation

Spleens from OT-I Tg mice were passed through a 100 μm cell filter and washed in PBS. The splenocytes were resuspended in 5ml ACK Lysis Buffer (0.1 mM EDTA, 150 mM NH_4_Cl from Sigma, 10 mM KHCO_3_ from Fisher Scientific) and incubated for 5 minutes at RT. ACK Lysis Buffer was then removed by centrifuging the cells at 450 g for 5 minutes, and the supernatant removed. Splenocytes were then washed in PBS by resuspending in 10 ml PBS and centrifuging at 450 g for 5 mins before removing the supernatant. If splenocytes required Carboxyfluorescein succinimidyl ester (CFSE, ThermoFisher)-labelling, this was performed by incubating the cells at 5×10^7^ cells/ml in 10 μM CFSE diluted in PBS-0.1% BSA for 6 mins at 37 °C, before washing twice with 25 ml cRPMI. Control, hB7-H3, mCD70 or m4-1BBL MC38 cells were harvested from culture and incubated at 1×10^6^ cells/ml with 1 nM OVAp for 30 minutes in a 37 °C shaking incubator. After washed with 10 ml of cRPMI, the MC38 cells were irradiated with 100 Gy using an X-ray source (350 kV, 11.4 mA, SSD: 29.5, Field Size Diameter: 20.4 cm, Filter: Al 2.00 mm) prior to resuspending in cRPMI. Splenocytes (1×10^5^) and MC38 cells (1×10^3^) were co-cultured in a 96-well flat-bottomed plate in 200 µl and, in some cases, specified bsAb were added. The co-culture was incubated at 37 °C for either two days to assess CD25 expression or three days, for all other conditions. Gating strategies shown in Supplementary Figure 8.

### Human T cell proliferation

Human PBMCs were isolated from anonymised leukocyte cones from healthy adult donors obtained through the NHS blood and transplant service. The use of human tissue was approved by the East of Scotland Research Ethics Service, Tayside, UK, and via the Faculty of Medicine Research Ethics Committee under submission 19660. PBMCs were isolated from leukocyte cones by density gradient centrifugation using Lymphoprep (Stemcell, 18060). Isolated PBMCs were labeled with 2 μM CFSE (ThermoFisher Scientific, C34554) for 10 min. CFSE-labeled PBMCs (1×10^5^) were cultured with 1×10^4^ irradiated (100 Gy: 350 kV, 11.4 mA, SSD: 29.5, Field Size Diameter: 20.4 cm, Filter: Al 2.00 mm) control MC38 or MC38 hB7-H3 cells in a 96 well plate. PBMCs were stimulated with sub-optimal anti-CD3 (OKT3) 0.03 or 0.1 ng/ml and 0.1-10 nM of either isotype hIgG1, isotype bsAb, Urelumab bsAb, Utomilumab bsAb, hCD27.15 bsAb or Varlilumab bsAb. CFSE dilution in CD8^+^ T cells, as well as CD25 and Granzyme B expression, were measured after 72 hours by flow cytometry. CFSE low cells were identified by comparison to CFSE-labeled cells that were cultured in the absence of any stimulation. Gating strategy shown in Supplementary Figure 9.

### ELISA for IL-2

Capture and detection antibodies for IL-2 were purchased from BD Pharmingen (anti-mouse IL-2 (554424) and biotin-labelled anti-mouse IL-2 (554426). Capture antibody (1 µg/ml) was coated onto Nunc Maxisorp flat bottomed plates (ThermoFischer Scientific, 439454) overnight at 4 °C. The coating antibody was removed and the plate blocked with PBS + 1 % w/v BSA for 1 hour at RT. Supernatant (50 µL) from mouse OT-I CD8^+^ T cell proliferation assays was then added and incubated for 1.5 hours at RT. Following 3 washes with PBS + 0.05 % v/v Tween20, 1 µg/mL of detection antibody was added for 1 hour at RT. Following another 3 washes with PBS + 0.05 % v/v Tween20, detection antibody binding was visualised using high-sensitivity HRP-conjugated streptavidin (1:1000 dilution) and OPD substrate.

### Flow cytometry

Fluorescently labelled antibodies were purchased from eBioscience: allophycocyanin (APC)-labeled, APC-Cyanine (Cy)-7-labeled, Fluorescein isothiocyanate (FITC)-labelled, or eFluor (eF) 506-labeled anti-mCD8α (53-6.7), R-phycoerythrin (PE)-labelled anti-mCD25 (PC61.5), PE-labelled anti-mCD4 (RM4-5), PE-labelled anti-mOX40 (OX86), PE-labelled anti-mGITR (DTA-1), PE-labelled anti-m4-1BB (17B5), PE-labelled anti-eomesodermin (Dan11mag), PE-labelled anti-IFN-γ (XMG1.2), eF450-labeled anti-T-Bet (4B10), eF450-labelled CD45.2 (104), FITC-labelled anti-TNF-α (XT22); or Biolegend: PE-Cyanine 7-labeled anti-hCD8α (RPA-T8), Alexa Fluor 647-labelled anti-Granzyme B (GB11), PE-labelled anti-hCD25 (BC96); or BD Pharmingen PE-labelled anti-mCD27 (LG.3A10). For flow cytometry staining, if samples contained red blood cells, these were lysed using ACK buffer. Cells were then viability stained with either Fixable Viability Dye eF780 (eBioscience) or Fixable Viability Dye eF506 (eBioscience) as per the manufacturers’ instructions. FcγR-blocking was performed using either 10 µg/ml anti-FcγRII/III (clone: 2.4G2, produced in house) for mouse cells, or 10 % v/v human AB serum for human cells. This was performed at 4 °C 10 mins prior to incubation with primary antibodies for 30 mins at 4 °C. For intracellular staining, cells were fixed and permeabilised using the FoxP3/Transcription Factor Staining Kit, as per the manufacturers’ instructions. Cells were washed before analysis on a BD FACS Canto II or a BD FACS Fortessa using the BD FACSDiva software.

### Surface plasmon resonance (SPR)

Binding affinities of bsAb for their respective target receptors was determined by SPR using a Biacore T200 (Biacore). In brief, the target recombinant protein was captured using either an anti-hFc (Human Antibody Capture Kit; Cytiva, BR100839) or anti-His antibody (His Capture Kit; Cytiva, 28995056) that had been amine-coupled to a CM5 chip (Cytiva, 29104988), as per the manufacturer’s instructions. Recombinant proteins were captured at a concentration of 2.5 µg/ml, using a flow rate of 10 µl/min for 30 sec. BsAb were then diluted in HBS-EP+ (Cytiva, BR100669) and injected into the flow cells at decreasing concentrations at a flow rate of 30 μ/min, allowing 300 s for association and 300 s for dissociation. Sensor surfaces were regenerated between runs using either 3M MgCl_2_ for 1 minute at 20 µl/min (anti-hFc) or 10 mM Glycine-HCl pH1.5 for 1 minute at 30 µl/min (anti-His). Binding responses were reference-subtracted using a control flow cell, and kinetic parameters were determined by fitting the data to a Bivalent analyte model using the Biacore Evaluation Software (Cytiva). *K*_D_ values were calculated as *k*_d_/*k*_a_.

### Statistical analysis

GraphPad Prism 10 was used for statistical analysis. Ordinary one-way analysis of variance (ANOVA) with Tukey’s post-hoc multiple comparison test or a paired-t test was used as indicated in the figure legends throughout. Where error bars are shown, they indicate SEM as detailed in the figure legends. Tumour area was analysed by calculating the area under the curve (AUC) before performing a one-way ANOVA. Survival analysis was performed using log-rank (Mantel–Cox) test between indicated groups. *p<0.05, **p<0.01, ***p<0.001, **** p<0.0001.

## Acknowledgements

The authors thank Christine Penfold, Ian Mockridge and members of the Biomedical Research Facility, University of Southampton, for their help with antibody production and in vivo studies. The study was funded by Cancer Research UK Award number DRCDDRPGMApr2020\100005 (A.Al-S., S.A.B., M.S.C.).

