## Supplementary Table I and Figures for "Bispecific Antibody Architecture and TNFRSF Target Selection Determine CD8^+^ T Cell Differentiation and Anti-tumour Immunity"

### Supplementary information

| bsAb | TNFRSF $K_D$ (M) | B7-H3 $K_D$ (M) |
|---|---|---|
| 2x2 LOB12 m4-1BB x B7-H3 | $3.3 \times 10^{-9}$ | $2.2 \times 10^{-8}$ |
| 2x1 LOB12 m4-1BB x B7-H3 | $2.8 \times 10^{-9}$ | $7.4 \times 10^{-7}$ |
| 1x1 LOB12 m4-1BB x B7-H3 | $2.3 \times 10^{-8}$ | $1.5 \times 10^{-7}$ |
| 2x2'light' LOB12 m4-1BB x B7-H3 | $2.2 \times 10^{-9}$ | $1.1 \times 10^{-7}$ |
| 2x2'light' G4S LOB12 m4-1BB x B7-H3 | $2.1 \times 10^{-9}$ | $1.9 \times 10^{-8}$ |
| 2x2 AT124 mCD27 x B7-H3 | $1.6 \times 10^{-8}$ | $2.3 \times 10^{-8}$ |
| 2x2 DTA-1 mGITR x B7-H3 | $0.8 \times 10^{-9}$ | $2.7 \times 10^{-8}$ |
| 2x2 MRC OX86 mOX40 x B7-H3 | $1.4 \times 10^{-8}$ | $2.8 \times 10^{-8}$ |
| 2x2 Utomilumab x B7-H3 | $8.3 \times 10^{-10}$ | $7.5 \times 10^{-9}$ |
| 2x2 Urelumab x B7-H3 | $6.2 \times 10^{-10}$ | $8.0 \times 10^{-9}$ |
| 2x2 Varlilumab x B7-H3 | $1.6 \times 10^{-9}$ | $1.3 \times 10^{-8}$ |
| 2x2 hCD27.15 x B7-H3 | $6.0 \times 10^{-10}$ | $1.4 \times 10^{-8}$ |

**Supplementary Table 1**. **Binding affinities of bsAb to TNFRSF and B7-H3 proteins**. Kinetic parameters were determined by fitting SPR data to a Bivalent analyte model using the Biacore Evaluation Software. $K_D$ values were calculated as kd/ka.

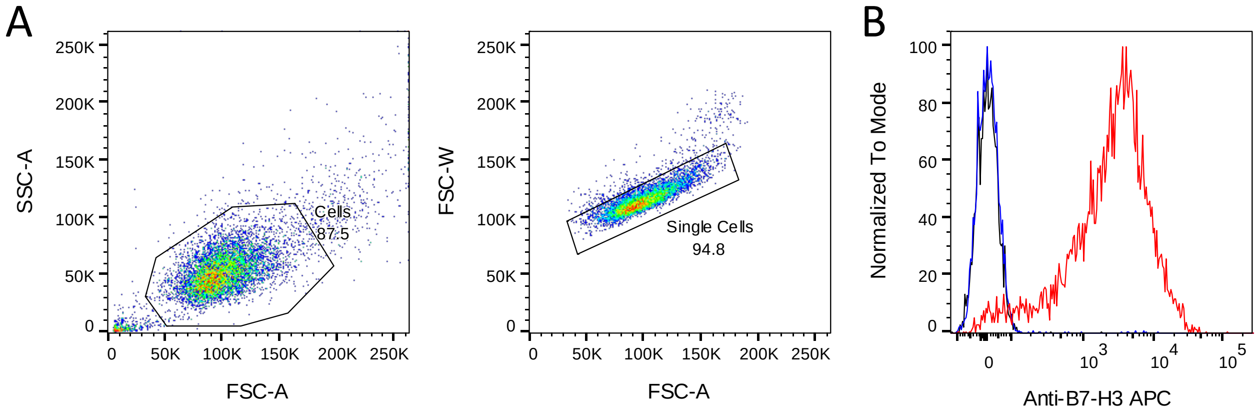


**Supplementary Figure 1. B7-H3 expression on MC38 hB7-H3⁺ cells.**
**a** Gating strategy and **b** expression of human B7-H3 on MC38 cells (blue) and hB7-H3⁺ MC38 cells (red), compared to an isotype control (black).

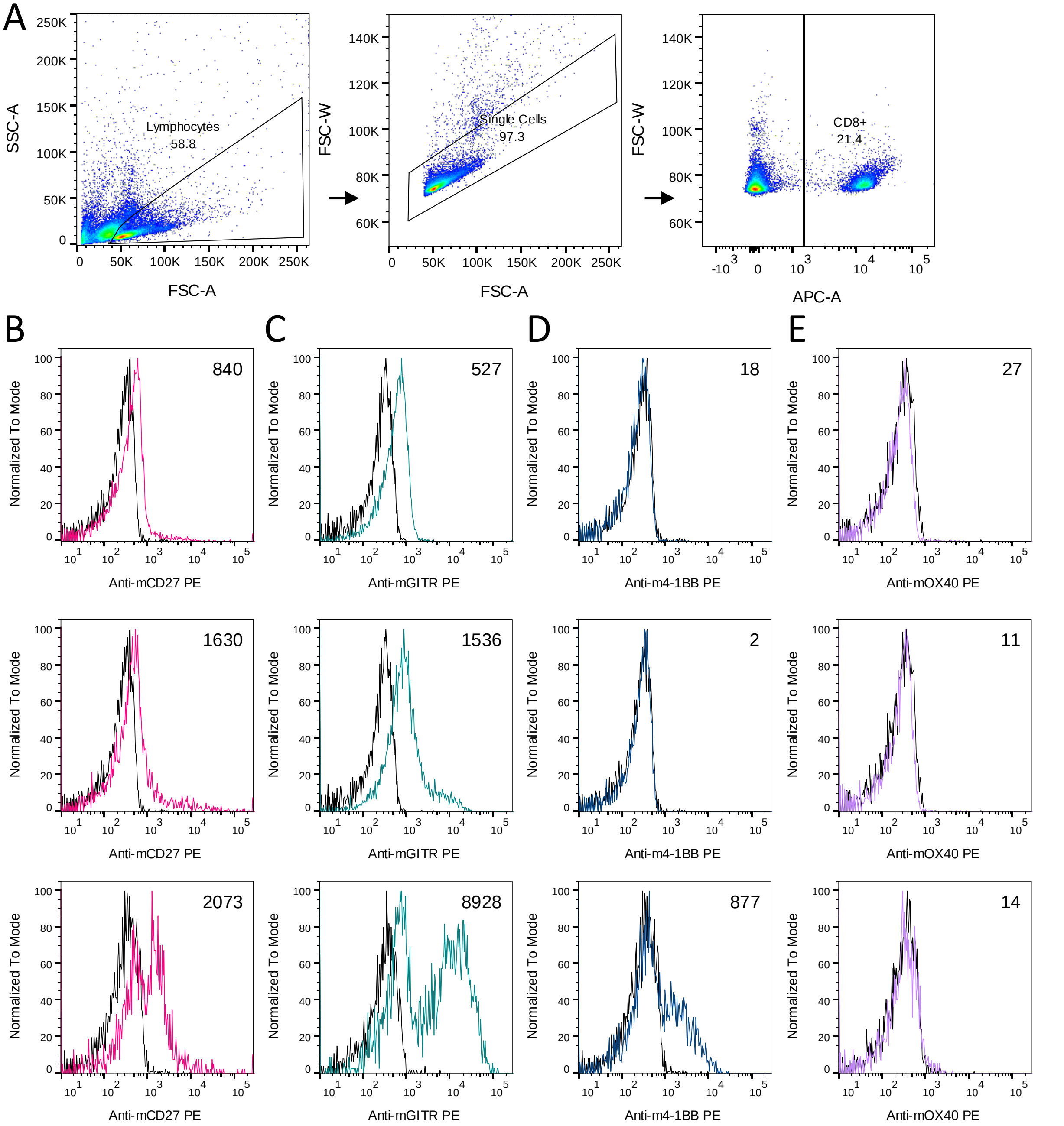


**Supplementary figure 2: TNFRSF expression on OT-I CD8⁺ T cells during stimulation.**
**a** Flow cytometry gating strategy. **b-e** Representative histograms showing **b** CD27, **c** GITR, **d** 4-1BB and **e** OX40 expression on OT-I CD8⁺ T cells following stimulation with OVAp-pulsed MC38 hB7-H3 cells after 0 hours (upper), 24 hours (centre) and 48 hours (lower). Inset number shows the MFI difference between staining antibody (colour) and isotype control (black). Data representative of 2 independent experiments.

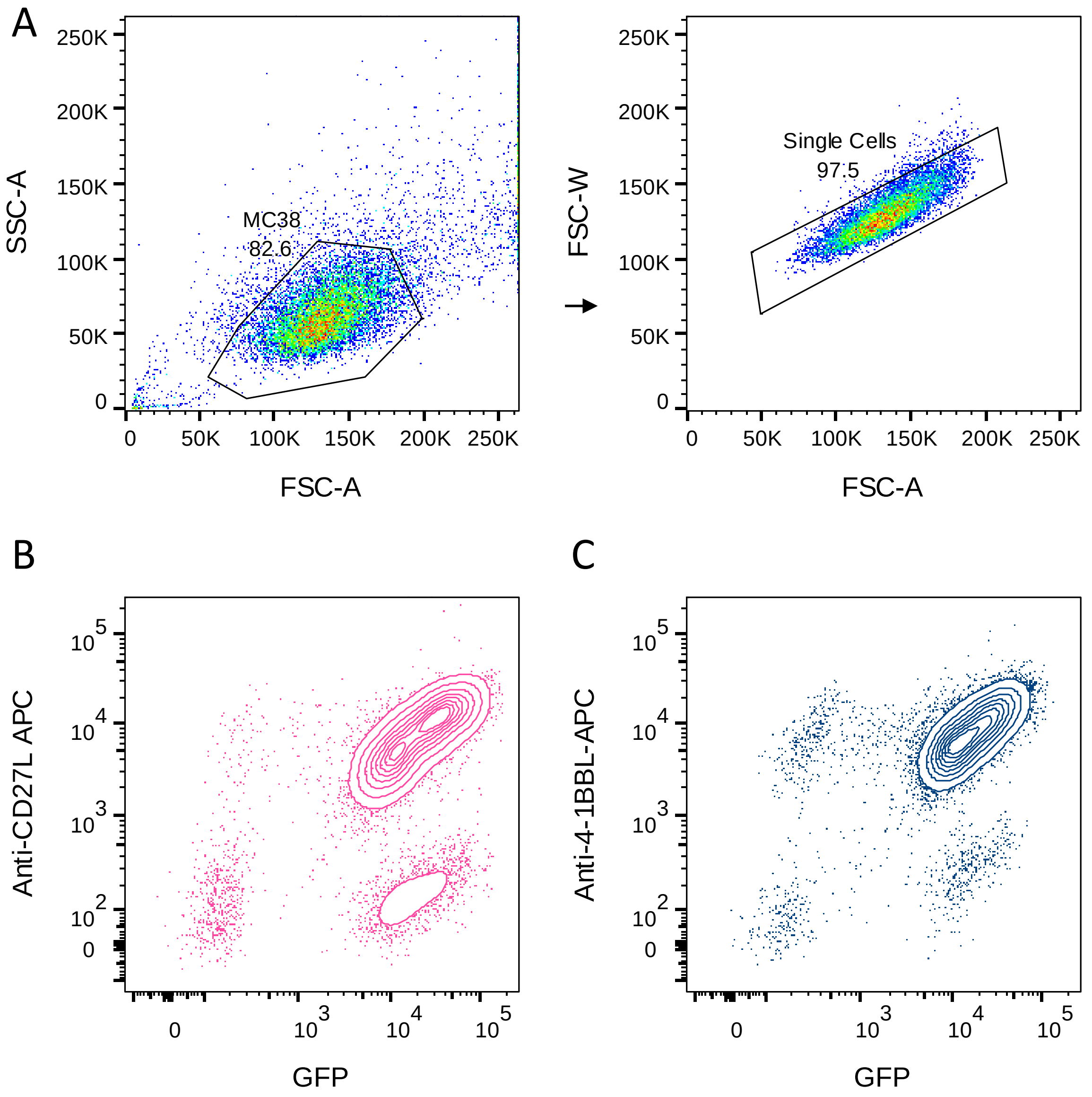


**Supplementary figure 3. TNFRSF ligand expression on MC38 B7-H3⁺ cells.**
**a** Flow cytometry gating strategy for MC38 cells expressing TNFRSF ligands. **b-c**. Contour plots of TNFSF expression on transduced hB7-H3⁺ MC38 cells. **b** CD27L and GFP expression on MC38 CD27L cells. **c** 4-1BBL and GFP expression on MC38 4-1BBL cells.

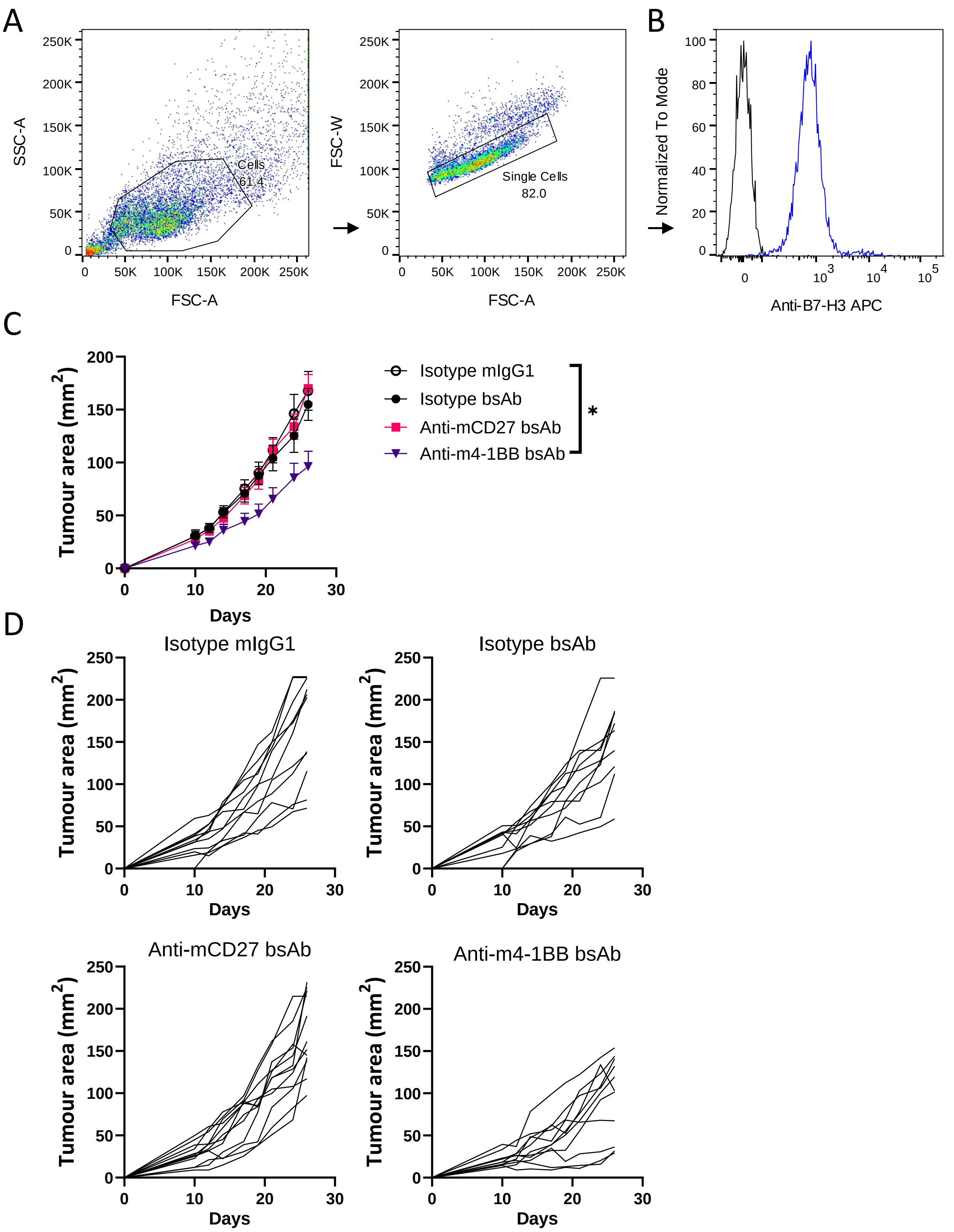


**Supplementary figure 4. Anti-tumour activity of TNFRSF-targeted bsAbs.**
**a** Flow cytometry gating strategy and **b** Expression of endogenous B7-H3 on 4T1 cells (blue) compared to an isotype control (black). **c-d** Tumour area of mice inoculated with $2.5 \times 10^4$ 4T1 cells following treatment with the specified antibodies on days 3, 5 and 7. Data points show **c** mean tumour area ± SEM or **d** individual tumour growth curves of 10-12 mice per group from two independent experiments. Statistical significance was determined by one-way ANOVA with multiple comparisons performed on AUC. *p<0.05.

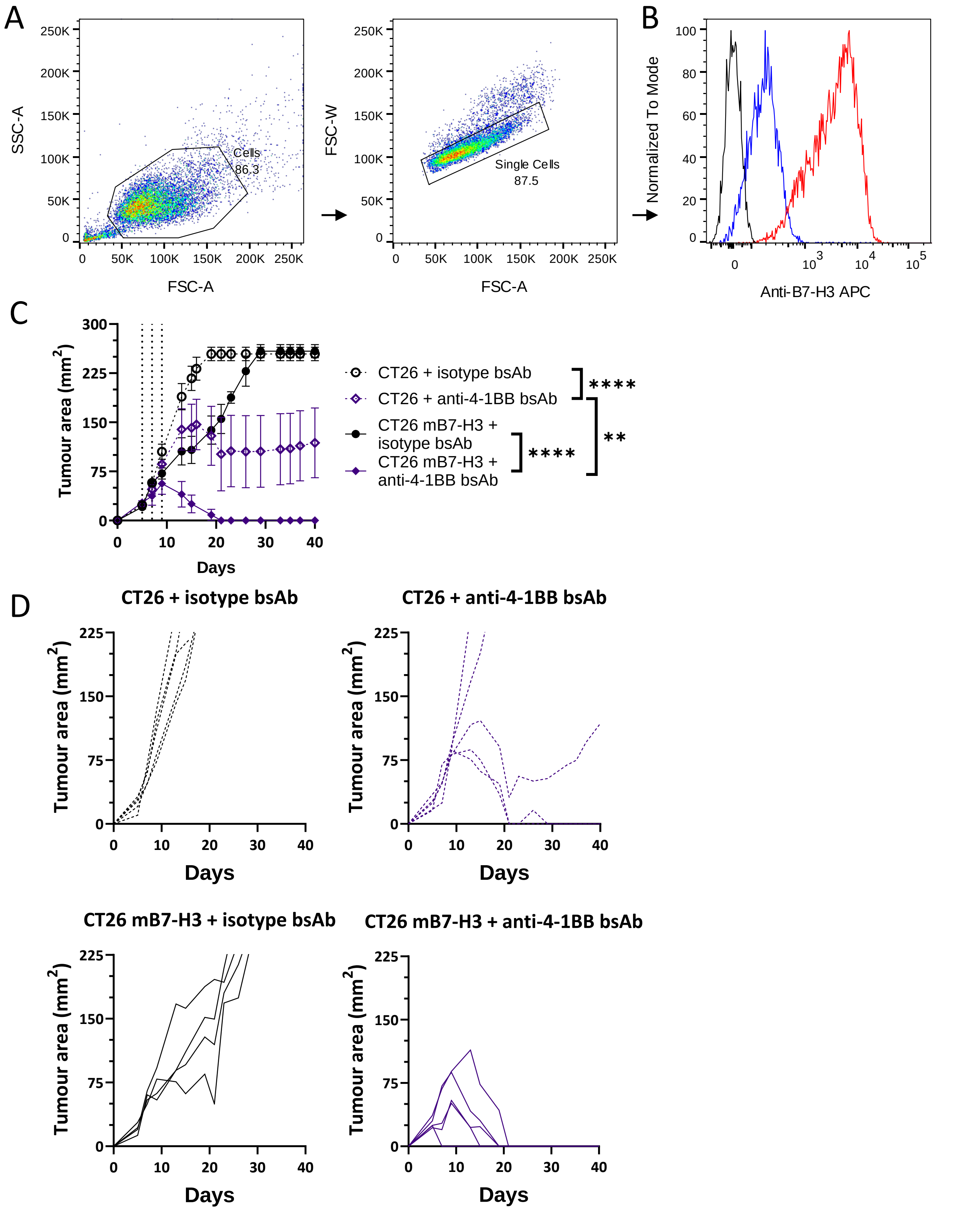


**Supplementary figure 5. B7-H3 density on CT26 impacts anti-4-1BB bsAb efficacy.**
**a** Flow cytometry gating strategy and **b** Expression of murine B7-H3 on CT26 cells (blue) and engineered CT26 to increase murine B7-H3 expression (CT26 mB7-H3 cells; red), compared to an isotype control (black). **c-d** Tumour area of Balb/c mice inoculated with $5x10^5$ control empty vector transduced CT26 cells (hollow, dashed lines) or CT26 mB7-H3 (filled, solid lines) cells following treatment with the specified antibodies on days 5, 7 and 9. Data points show **c** mean tumour area ± SEM or **d** individual tumour growth curves of 4-5 mice per group. Statistical significance was determined by one-way ANOVA with multiple comparisons performed on AUC. **p<0.01, ****p<0.0001.

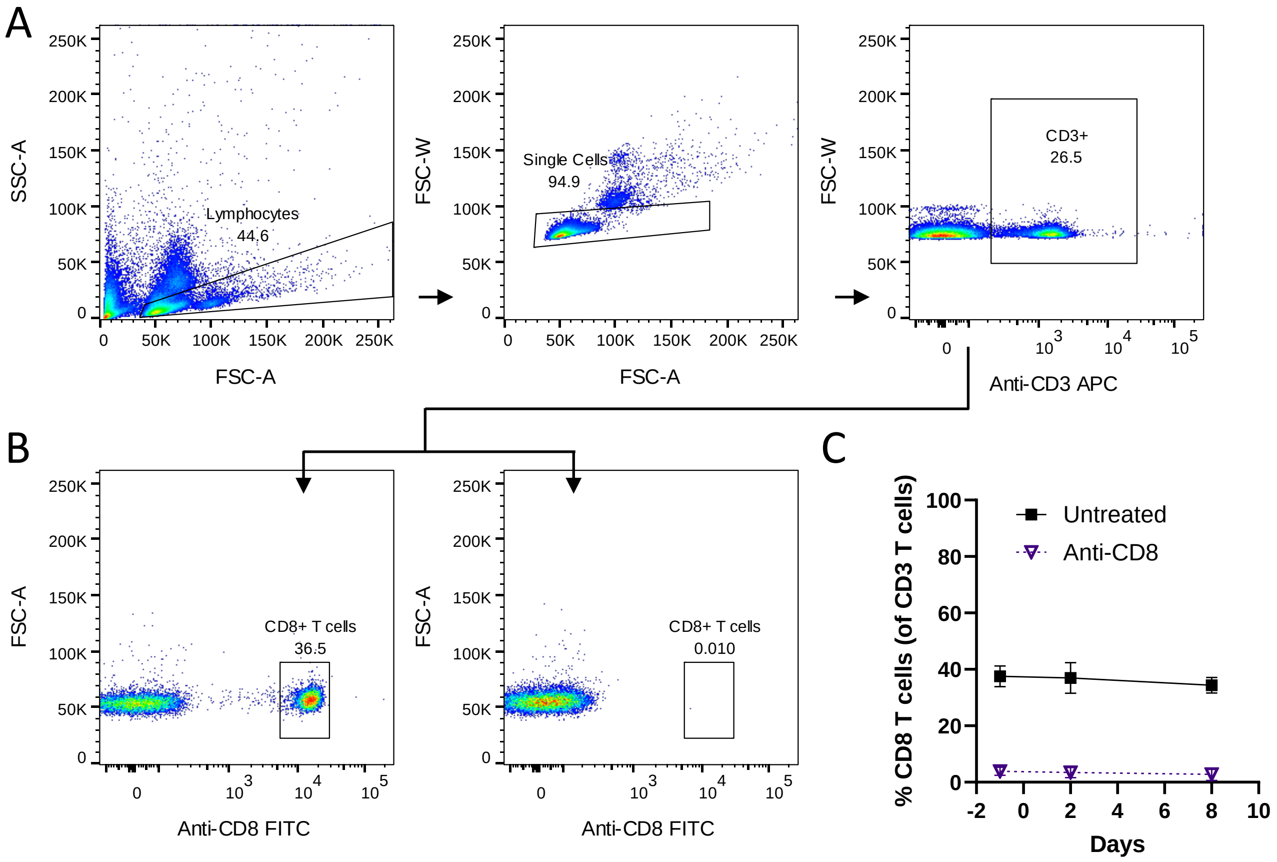


**Supplementary figure 6. Confirmation of CD8⁺ T cell depletion after anti-CD8 antibody injection.** C57BL/6 mice were inoculated s.c. with 5x10$^5$ MC38 hB7-H3 cells. To assess the role of CD8⁺ T cells, one group was injected i.p. with anti-CD8 antibody 2 days before and 1 day after tumour injection. **a-c. a** Flow cytometry gating strategy for detection of CD8⁺ T cells using non-competing anti-CD8 antibody , **b** representative dot plots demonstrating normal levels of CD8⁺ T cells (left) in control mice or CD8⁺ T cell loss in mice that received depleting an anti-CD8 antibody (right), and **c** time course of peripheral CD8⁺ T cell depletion. Data points show mean ± SEM of 6 mice per group.

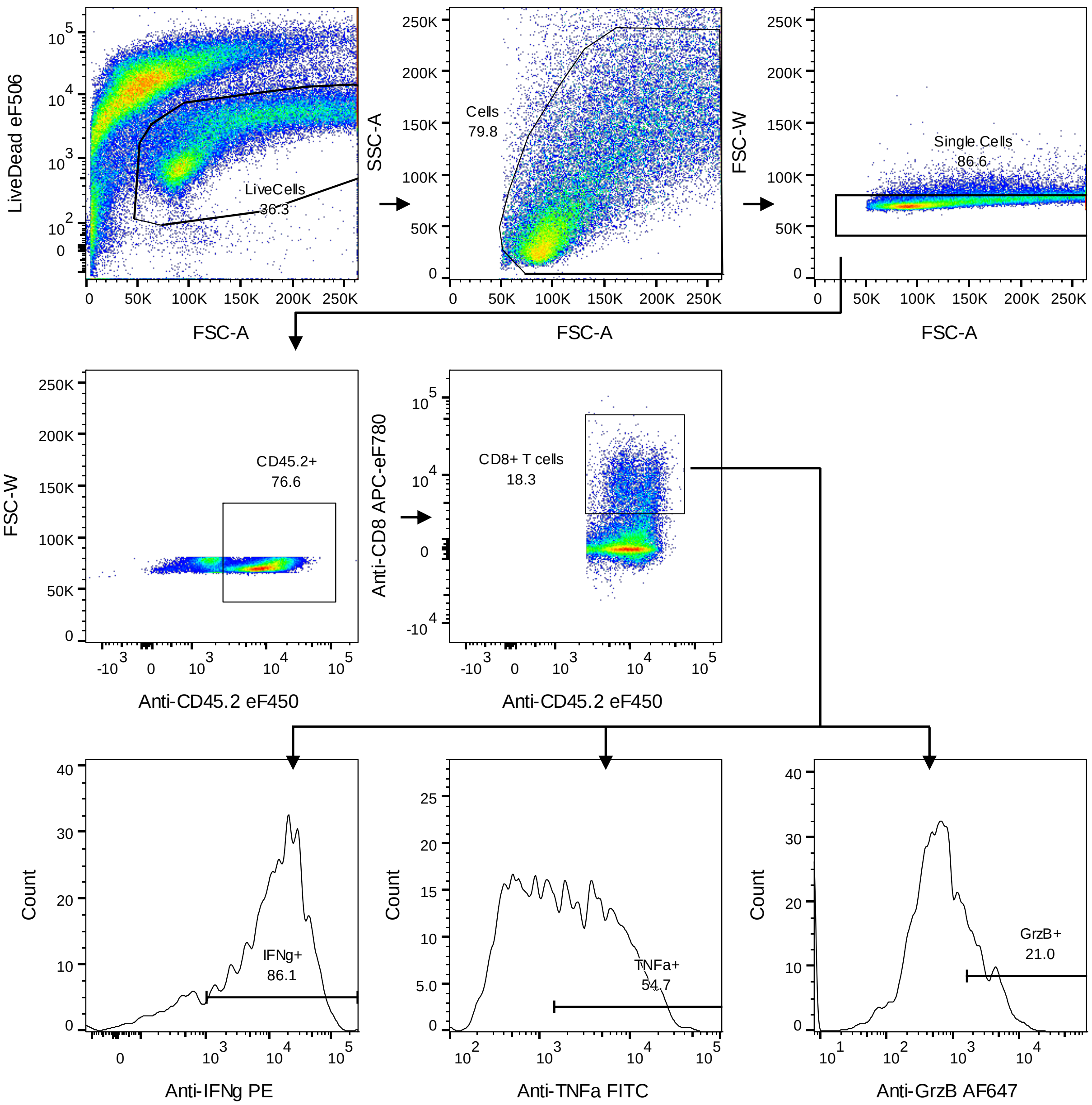


**Supplementary figure 7. Gating strategy for tumour infiltrating lymphocytes.**
Gating strategy to identify mouse CD8$^{+}$ T cells that have infiltrated MC38 tumours and the expression of IFNγ, TNFα and granzyme B (GrzB). Electronic gating in histogram plots was set based on isotype control antibody staining.

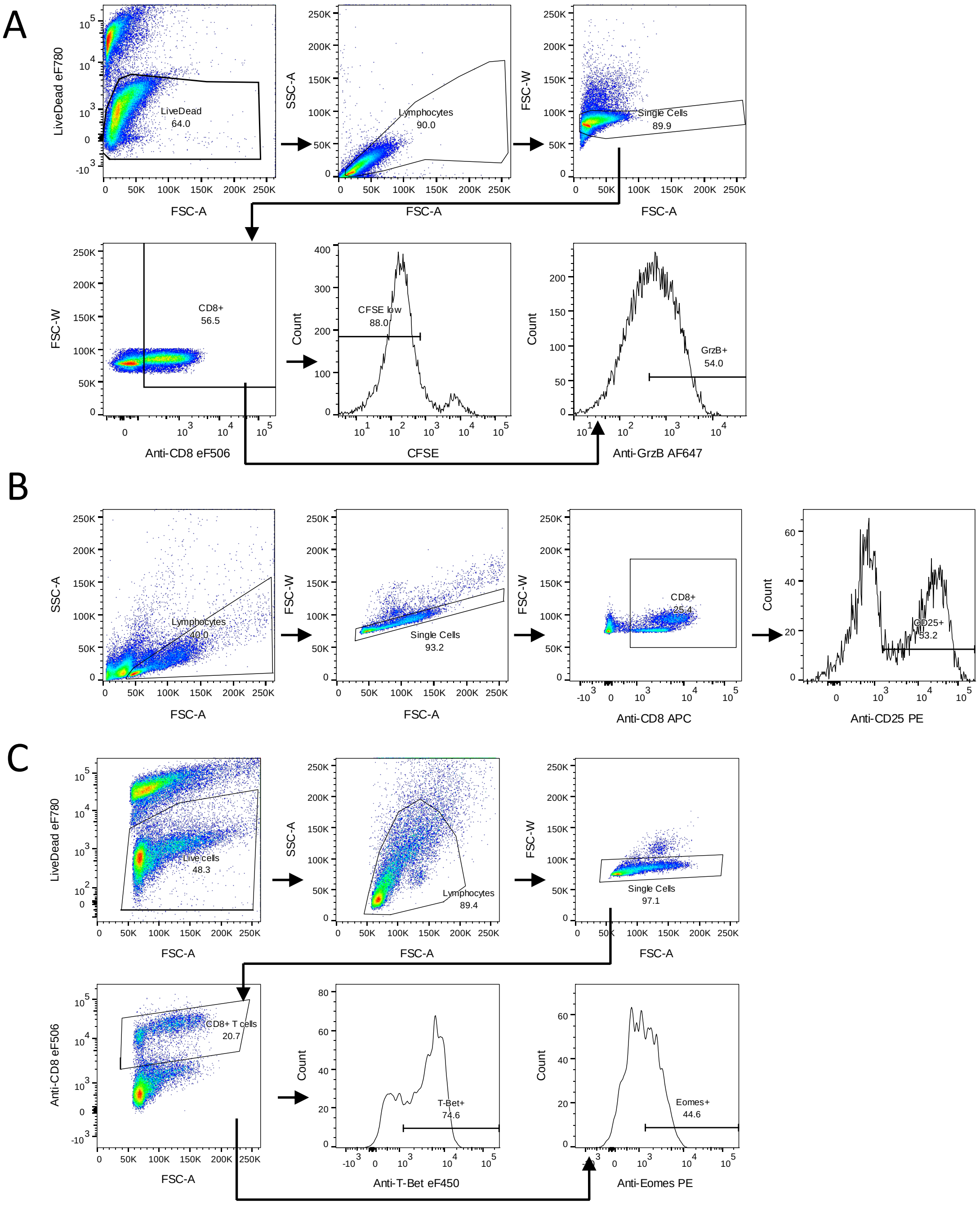


**Supplementary figure 8. Gating strategies for mouse CD8⁺ T cells.**

**a** Gating strategy to identify mouse CFSE low CD8⁺ T cells and granzyme B high CD8⁺ T cells. **b** Gating strategy to identify mouse CD25⁺ CD8⁺ T cells. **c** Gating strategy to investigate T-bet and Eomes expression in mouse CD8⁺ T cells. Electronic gating in histogram plots was set based on unstimulated CFSE high CD8⁺ T cells or isotype control antibody staining.

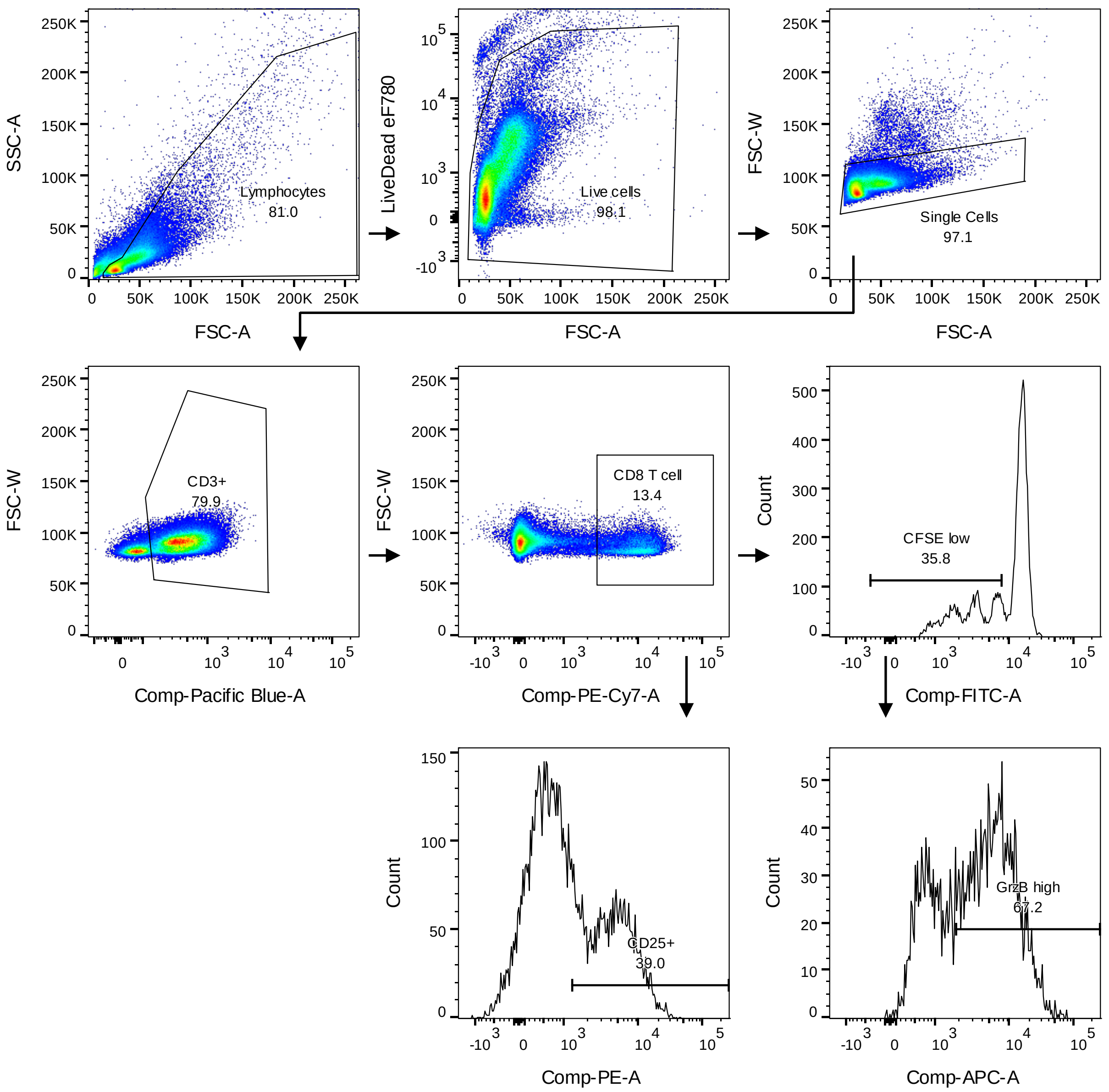


**Supplementary figure 9. Gating strategy for human CD8⁺ T cells.**
Gating strategy to identify human CFSE low CD8⁺ T cells, CD25⁺ CD8⁺ T cells, and CFSE low Grz B high CD8⁺ T cells. Electronic gating in histogram plots was set based on unstimulated CFSE high CD8⁺ T cells or isotype control antibody staining.
